# Chromogranin A regulates the dynamics of neurosecretion through its interaction with phosphatidic acid

**DOI:** 10.64898/2026.01.29.699889

**Authors:** Thomas Ferrand, Alexander Wolf, Fanny Laguerre, Lina Riachy, Antoine Schlichter, Lydie Jeandel, Magalie Bénard, Brice Beauvais, Damien Schapman, Sylvette Chasserot-Golaz, Cathy Royer, Caroline Bérard, Dorthe Cartier, Jiyoung Lee, Luca Grumlolato, Ludovic Galas, Nicolas Chartrel, Pierre-Yves Renard, Youssef Anouar, Sébastien Balieu, Nicolas Vitale, Maité Montero-Hadjadje

## Abstract

Altered neurosecretion is a common feature of diverse pathophysiological conditions, including central nervous system disorders, hypertension, and tumorigenesis, where chromogranin A (CgA) is widely used as a biomarker, and both phosphatidic acid (PA) synthesis and catecholamine (CA) secretion are dysregulated. Here, we identify a direct interaction between CgA and PA at the plasma membrane of living cells using a newly developed synthetic PA fluorescent probe and Förster’s resonance energy transfer (FRET) combined with fluorescence lifetime imaging microscopy (FLIM). Confocal microscopy and transmission electron microscopy (TEM) further reveal that this interaction is spatially confined to exocytic sites. Using Total Internal Reflection Fluorescence microscopy (TIRF-M), we show that expression of a CgA variant lacking the PA-binding domain (PABD) in COS-7 cells increases the frequency of exocytic events and accelerates CgA release kinetics. In chromaffin cells, amperometry and live tracking of exocytosis-endocytosis demonstrate that CgA overexpression enhances granular CA content, extends fusion pore opening, and accelerates exocytosis-endocytosis coupling, effects that are abolished upon expression of CgA variant. Together, these findings unveil a new role of CgA/PA interaction in fine-tuning neurohormone secretion, suggesting unexplored avenues for restoring neurosecretion in disease-relevant contexts.

## Introduction

Neurosecretion is frequently disrupted in neuroendocrine tumors and in metabolic or neurodegenerative diseases, reflecting alterations in hormone and neurotransmitter release from neurons and neuroendocrine cells. This process relies on the regulated secretory pathway (RSP), which governs the packaging and stimulus-dependent release of hormones, neuropeptides, and neurotransmitters to sustain intercellular communication and numerous physiological functions. In neuroendocrine cells, the RSP is orchestrated by dense-core secretory granules (SG) that bud from the *trans*-Golgi network (TGN) and then accumulate in large numbers within the cytoplasm, where they support rapid, activity-dependent secretion.

Chromaffin cells of the adrenal medulla have served as a central model to establish the molecular mechanisms underlying regulated exocytosis^1^. Their SG are enriched in catecholamines (CA), granins such as chromogranin A (CgA), neuropeptides including neuropeptide Y, ATP, and divalent cations. Circulating CgA levels are increased in several pathologies and its expression is commonly upregulated in neuroendocrine tumors^2^, establishing CgA as a clinically used biomarker^3^. However, this also raises the question of whether this glycoprotein plays roles beyond disease association.

Functionally, CgA is a multifaceted protein. It interacts with SG components to stabilize the dense-core matrix, it is proteolytically processed into bioactive peptides with endocrine, paracrine, and autocrine actions, while also contributing to SG biogenesis at the TGN^4,5^. Remarkably, expression of CgA in non-endocrine COS-7 cells, lacking endogenous SG, is sufficient to drive the formation of SG-like organelles in a manner dependent on its terminal regions^6,7^. Studies in neuroendocrine cells have underscored the importance of these domains, whose helical motifs are thought to favor interactions with biological membranes^7,8^.

Among these structural regions, we recently identified a dedicated phosphatidic acid (PA)-binding domain (PABD) within CgA that may participate in the remodeling of TGN membranes^9^. Elevated local PA concentrations, a conical lipid known to promote membrane negative curvature, have been implicated in shaping chromaffin cell membranes^10,11^. We therefore proposed that PA clustering, driven by CgA accumulation at specific TGN sites, could generate the membrane curvature required to initiate SG budding and maturation^9^.

Resulting SG are routed to the plasma membrane, where they are sequestered within the cortical actin network. Under stressful conditions, sympathetic stimulation increases cytosolic Ca^2+^, triggering docking and fusion of SG and resulting in partial or complete discharge of their cargo. Depending on physiological demand and the extent of Ca^2+^ influx, regulated exocytosis can proceed through at least three distinct modes^12,13^. In the “kiss and run” mode, a transient fusion pore opens, allowing selective release of small molecules such as CA while retaining macromolecules within the SG. The “kiss and stay” mode supports partial release of neuropeptides while maintaining the characteristic omega-shaped granule profile. In contrast, “full fusion” enables complete emptying of SG content and flattening of the SG membrane upon merger with the plasma membrane^14^. Despite extensive work, the molecular control of these exocytic modes and their transitions remains incompletely understood.

CgA has also emerged as a contributor to late stages of neurosecretion. Indeed, it interacts with the Soluble N-ethylmaleimide-sensitive-factor Attachment protein REceptor (SNARE) protein syntaxin 1A and the calcium sensor synaptotagmin-1, both core components of the minimal fusion machinery governing regulated exocytosis^15^. In addition, interactions between CgA and CA within the SG lumen have been demonstrated to modulate SG matrix expansion, therefore also tuning exocytosis^16,17^. Moreover, CgA has also been shown to control fusion pore expansion during regulated secretion in chromaffin cells^18^. More recently, we demonstrated that CgA specifically binds specific monounsaturated and polyunsaturated PA species^9^, which themselves have been reported to regulate SG docking and fusion pore dynamics, respectively^19^. Together, these observations suggest that, through its capacity to interact with membrane PA and reshape membrane topology, as well as its engagement with the fusion machinery, CgA may act as a modulator of fusion pore stability and dynamics, thereby fine-tuning SG content release and their mode of exocytosis.

To evaluate this hypothesis, we used fluorescence lifetime imaging microscopy (FLIM) combined with Förster’s resonance energy transfer (FRET) to monitor the interaction between CgA and PA at the plasma membrane of living cells. With the advent of highly sensitive and photon-efficient FLIM instrumentation, this approach has become particularly suited to detecting fast changes in FRET that report transient molecular interactions. Such dynamics are characteristic of regulated exocytosis in neuroendocrine cells, where SG fusion occurs rapidly upon stimulation and SG membranes are subsequently retrieved to preserve plasma membrane homeostasis. Using this strategy, we found that CgA interacts with PA through its PABD at exocytic sites in chromaffin cells. Total Internal Reflection Fluorescence microscopy (TIRF-M) experiments further revealed that CgA/PA interaction shapes the dynamics of CgA secretion in COS-7 cells. Moreover, carbon fiber amperometry demonstrated that this interaction controls CA release kinetics and fusion pore expansion in primary chromaffin cells. Finally, confocal imaging showed that it alters the speed of SG membrane reuptake, thereby potentially regulating exocytosis-endocytosis coupling. Collectively, these findings establish CgA/PA interaction as a key determinant in the fine control of neurosecretion and suggest that targeting this interaction may provide opportunities to modulate altered neurosecretion across diverse pathophysiological contexts.

## Results

### CgA interacts with PA at the plasma membrane during regulated exocytosis

In a previous study, we showed that CgA interacts with PA in non-cellular systems, and that this interaction contributes to TGN membrane remodeling and budding, ultimately promoting SG biogenesis in neuroendocrine cells^9^. Here, we used FLIM-FRET to probe this interaction at the plasma membrane of living COS-7 cells. To this end, we engineered plasmids encoding CgA or a CgA mutant lacking its PABD (CgAΔPABD) fused to mKate2 as the fluorescence donor, and combined them with a novel synthetic PA probe labeled with ATTO647N^20^, serving as a fluorescence acceptor. It is of note that this synthetic form of PA was recently extensively characterized and shown to fully recapitulate the function of natural PA^20^. The lifetime of mKate2 fluorescence was recorded in COS-7 cells overexpressing CgA- or CgAΔPABD-mKate2 following BaCl_2_ stimulation and incubation with PA-ATTO647N (**Figure 1a**). When donor and acceptor are in close proximity, FRET occurs and the donor fluorescence’s lifetime decreases (**Figure 1b**). Importantly, the FLIM-FRET approach relies solely on donor fluorescence lifetime, which remains independent of fluorophore concentration, thereby enabling accurate interaction quantification despite local fluctuations.

**Figure 1.**
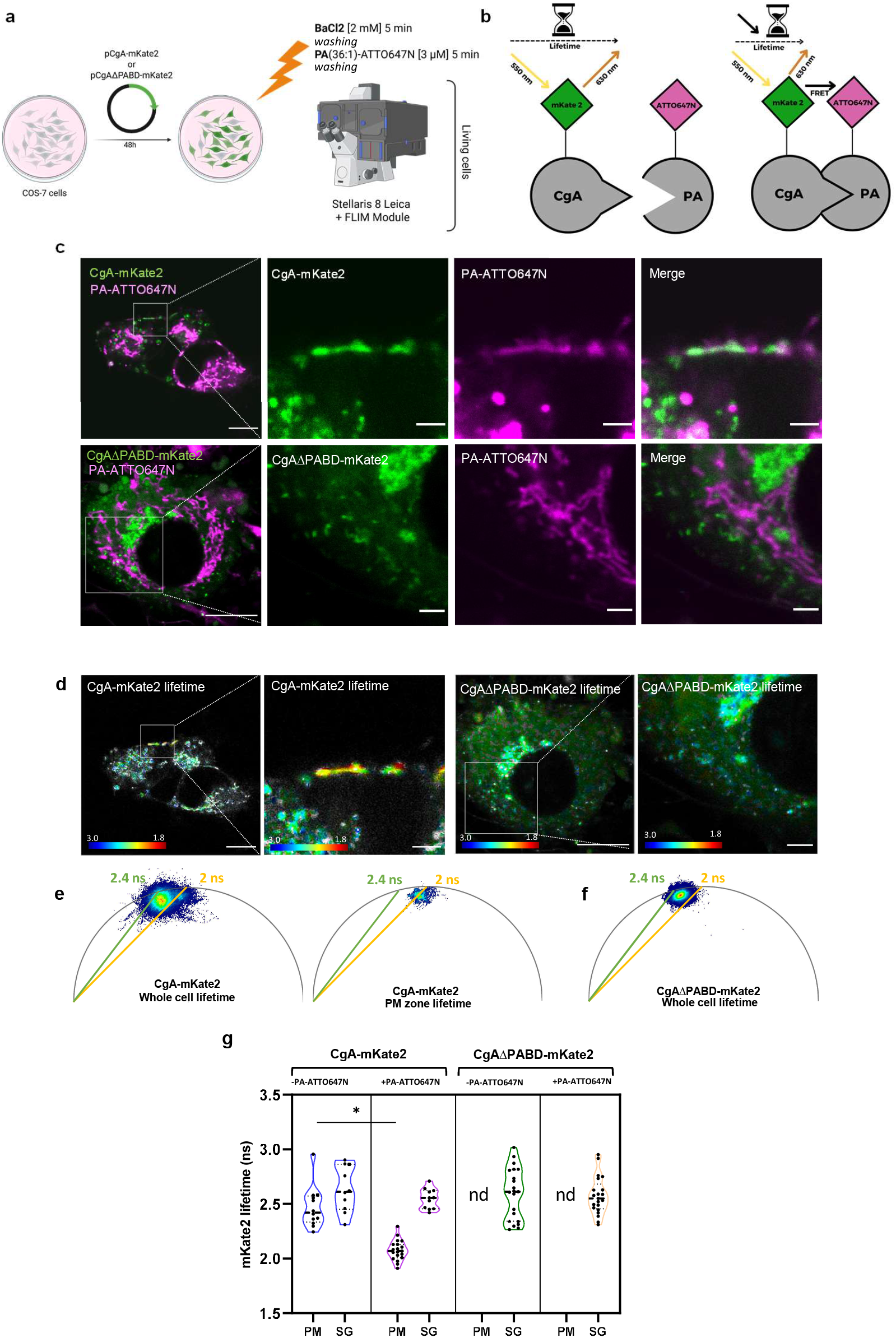
CgA interacts with PA at the plasma membrane after regulated exocytosis. (a) Schematic showing the experimental plan for CgA/PA interaction monitoring in living cells by the FLIM-FRET technique. Following the overexpression of CgA- or CgAΔPABD-mKate2 in COS-7 cells, their secretion was induced during 5 min using a BaCl_2_ solution. Then cells were washed and incubated 5 min with PA-ATTO647N. (b) Principle of FLIM-FRET technique to track the lifetime of the fluorescence donor (mKate2) without or with CgA/PA-ATTO647N interaction. The more CgA-mKate2 interacts with PA, the more the lifetime of mKate2 decreases due to its energy transfer to the acceptor fluorophore (ATTO647N). (c) Confocal observations of COS-7 cells overexpressing CgA-mKate2 or CgAΔPABD-mKate2 (green) after secretion stimulation and incubation with the PA-ATTO647N probe (magenta). Scale bars: 10 µm. The regions delimited by a white square on left images are enlarged to show the distribution of each fluorescent molecule. Scale bars: 2 µm. (d) Confocal observations of COS-7 cells overexpressing CgA-mKate2 or CgAΔPABD-mKate2, after cell incubation with PA-ATTO647N, showing the intensity of mKate2 fluorescence (grey), and mKate2 lifetime between 3 and 1.8 ns (rainbow color bar). The regions delimited by a white square are enlarged to show mKate2 lifetime at the plasma membrane. Scale bars: 5 µm. (e) Representative phasor plots showing the CgA-mKate2 lifetime at the level of the whole image and at the plasma membrane zone. Green line corresponds to 2.4 ns and yellow line to 2 ns, used as visual landmarks. (f) Representative phasor plot showing the CgAΔPABD-mKate2 lifetime at the level of the whole image. Green line corresponds to 2.4 ns and yellow line to 2 ns, used as visual landmarks. (g) Quantification of CgA-mKate2 or CgAΔPABD-mKate2 lifetime at the level of the plasma membrane (PM) or secretory granule (SG) without (-) or with (+) PA-ATTO647N. nd: not detected. Kruskall-Wallis test * p<0,05, each point representing the mean mKate2 lifetime in a ROI at the PM or SG level of 2 to 3 cells.

As a control, mKate2 lifetime in stimulated COS-7 cells expressing CgA- or CgAΔPABD-mKate2 was comparable prior to PA-ATTO647N addition (≈ 2.3 ns; **Supplemental Figure 1a, b**). Likewise, adding PA-ATTO647N to resting COS-7 cells expressing either construct did not alter donor lifetime (**Supplemental Figure 1c, d**). Following stimulation, PA-ATTO647N incorporated into both plasma membrane and endomembrane structures, but not into SG (**Figure 1c**), in agreement with previous observations^20^. Interestingly, the overall lifetime distribution differed between CgA- and CgAΔPABD-expressing cells (**Figure 1d, e, f**), owing to a significant lifetime decrease for CgA-mKate2 at the plasma membrane (**Figure 1d, e**). Because CgAΔPABD-mKate2 was not detected at the plasma membrane (**Figure 1c**), no lifetime change was measured in this compartment (**Figure 1d, f**).

Subcellular quantification across plasma membrane regions revealed a marked decrease in CgA-mKate2 lifetime following PA-ATTO647N addition (from 2.47 ± 0.19 ns to 2.08 ± 0.09 ns), indicating robust FRET and thus direct proximity between CgA and PA (**Figure 1g**). No significant lifetime modification was detected within SG (from 2.63 ± 0.21 ns to 2.55 ± 0.09 ns) (**Figure 1g**). Similarly, no change was detected for CgAΔPABD-mKate2 in SG (from 2.59 ± 0.24 ns to 2.58 ± 0.17 ns) (**Figure 1g**), and absence from the plasma membrane prevented lifetime assessment in that compartment. Together, these findings indicate that CgA interacts with PA through its PABD specifically at the plasma membrane during regulated exocytosis. To our knowledge, this is the first study to apply FLIM-FRET to monitor protein-lipid interactions in a dynamic context, as this method has been primarily used to probe protein-protein interactions^21^.

### CgA is localized at exocytic sites

Stimulation of SG exocytosis induces release of soluble cargos together with insertion of SG membrane proteins into the plasma membrane. Lumenal proteins thus transiently contact the extracellular environment upon fusion pore expansion. Using the approach described previously to visualize dopamine-β-hydroxylase (DBH) associated to SG membrane after regulated exocytosis^22^, primary cultured bovine chromaffin cells were stimulated during 5 min with 40 μM acetylcholine (ACh) in presence of anti-CgA antibody and subsequently fixed and permeabilized before secondary antibody application (**Figure 2a**). Confocal microscopy revealed a significant increase in CgA-labelled puncta, compared with unstimulated chromaffin cells (**Figure 2b**). This observation suggests that a fraction of intragranular CgA that was transiently accessible to anti-CgA antibodies remained associated to the cell surface after SG fusion and is recycled by compensatory endocytosis.

**Figure 2.**
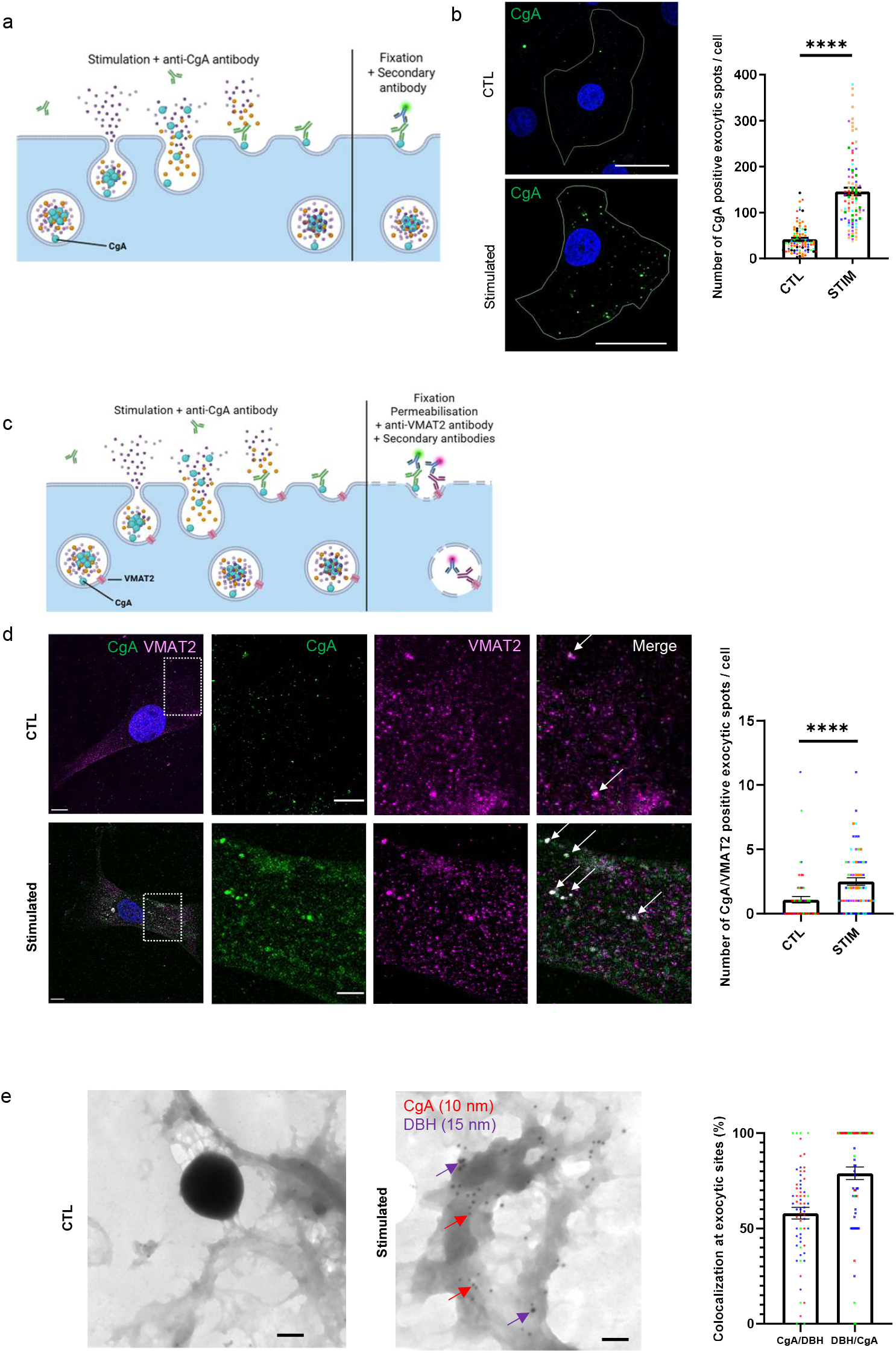
CgA is localized at exocytic sites after regulated secretion. (a) Schematic showing the experimental procedure for the incubation of living chromaffin cells with anti-CgA antibody, then fixed and processed for immunofluorescence with Alexa-488-conjugated secondary antibody to examine CgA at the plasma membrane. (b) Representative confocal images of chromaffin cells in resting condition (CTL) or stimulated (STIM) with acetylcholine (40 µM) for 10 min. Plasma membrane images are shown and were used to automatically quantify the number of CgA-positive exocytic sites. Values for this number are plotted as the means ± SEM (n=6, 90 cells per condition). Each point represents one analysed cell and are color coded according to experimental repeat. ***p < 0.0001, Mann-Whitney test. Scale bars: 10 µm. (c) Schematic showing the experimental procedure for the incubation of living chromaffin cells, in resting condition (CTL) or stimulated (STIM) conditions, with anti-CgA antibody, then fixed and processed for immunofluorescence with VMAT2 antibody. Anti-CgA antibody was revealed with Alexa-488-conjugated secondary antibody, and anti-VMAT2 antibody was revealed with Alexa-594-conjugated secondary antibody. (d) Observations using confocal microscopy of chromaffin cells in resting (CTL) or stimulated (STIM) conditions. White arrows show representative CgA and VMAT2 colocalized exocytic sites. Values for the number of CgA/VMAT2 positive exocytic sites are plotted as mean ± SEM (n=4 independent experiments with a total of 59 cells in resting condition and n=5 independent experiments with a total of 63 analyzed cells in stimulated condition). Each point is one analyzed cell and each color is an independent experiment. ****p < 0.0001, Mann-Whitney test. Scale bars: 10 µm. The regions delimited by a white square on left images are enlarged to show the distribution of each fluorescent molecule. Scale bars: 5 µm. (e) Electron microscopy of dual staining of CgA (10-nm gold beads, red arrows) and DBH (15-nm gold beads, purple arrows) on the outer face of the plasma membrane sheets obtained from nicotine-stimulated bovine chromaffin cells. Quantification of CgA/DBH and DBH/CgA colocalizations at exocytic sites are plotted as a percentage of the plasma membrane staining (65 cells analyzed from 3 independent experiments). Each point in the plot is one analyzed cell and each color is an independent experiment. Scale bars: 100 nm.

To further define their spatial organization, we performed a colocalization study with vesicular monoamine transporter 2 (VMAT2), a granular transmembrane protein (**Figure 2c**), and observed a significant increase of CgA colocalization with VMAT2-positive structures after cell stimulation consistent with exocytic sites (**Figure 2d**). These data indicate that cell surface-bound CgA arises from SG membrane fusion during regulated exocytosis.

We next examined the ultrastructural organization of CgA at the plasma membrane using immunogold labeling on plasma membrane sheets, as previously described^23,24^. Resting or nicotine-stimulated chromaffin cells were incubated with anti-CgA and anti-DBH antibodies to label fused SG membranes^22,25^, followed by secondary antibodies coupled to 10 nm and 15 nm gold particles, respectively. DBH positive clusters, corresponding to fused SG with the plasma membrane after cell stimulation, frequently contained CgA labelling (**Figure 2e**). Together these data showed the presence of CgA at exocytic sites after SG fusion in primary neuroendocrine cells, similar to DBH^22^.

### CgA/PA interaction regulates dynamics of CgA secretion

To investigate the functional consequences of CgA/PA interaction at exocytic sites, CgA secretion was monitored by TIRF-M in COS-7 cells overexpressing CgA- or CgAΔPABD-pHluorin and CgA- or CgAΔPABD-EGFP, before and after cell stimulation with a 2 mM BaCl_2_ solution. Because pHluorin fluorescence increases upon luminal pH neutralization, exocytic events could be readily tracked. While inducing a non-significant increase before stimulation (**Supplemental Figure 2**), overexpression of CgAΔPABD-pHluorin significantly increased the amount of regulated exocytosis events compared to CgA-pHluorin (from 25 ± 6 to 131 ± 35 events/cell/µm^2^/s x 10^6^) (**Figure 3a, b**) (**Videos 1** and **2**).

**Figure 3.**
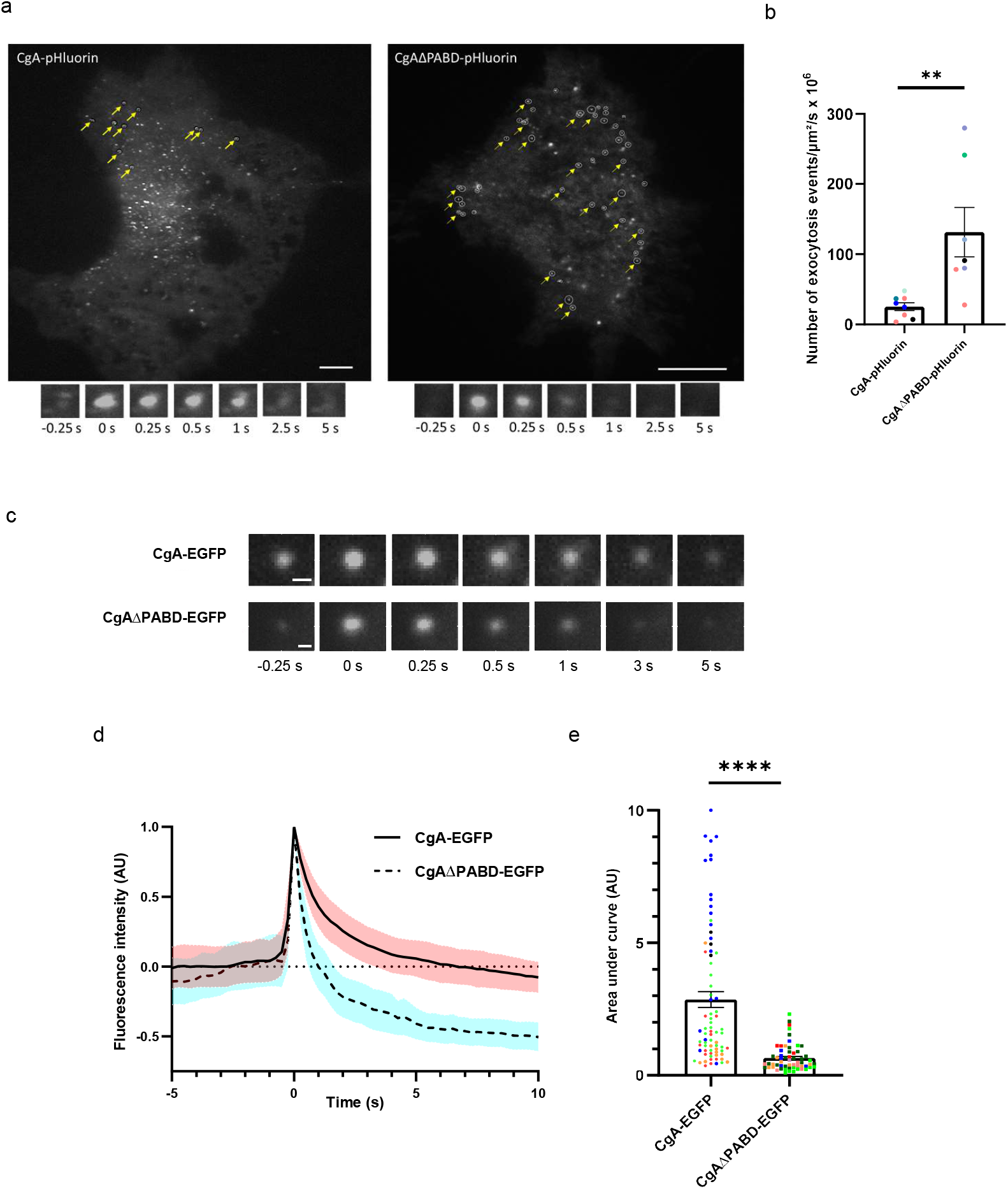
CgA interaction with PA through its PABD governs its regulated secretion. (a) TIRF-M imaging of COS-7 cells overexpressing and secreting CgA- and CgAΔPABD-pHluorin after 2 mM BaCl_2_ stimulation. SG leading to exocytic events on whole cells are surrounded and indicated by yellow arrows. Scale bars: 10 µm. Below are represented typical images of CgA- or CgAΔPABD-pHluorin exocytosis events. (b) Quantification of the number of exocytosis events in COS-7 cells overexpressing CgA-pHluorin and CgAΔPABD-pHluorin after their stimulation. (n=4 independent experiments; 8 cells in CgA condition and 7 cells in CgAΔPABD condition). Data are represented as mean ± SEM. Each point is one analyzed cell and each color is an independent experiment. **p < 0.01, Mann-Whitney test. (c) Typical images of CgA-EGFP or CgAΔPABD-EGFP exocytosis events. Scale bar: 500 nm. (d) Curves represent the normalized variation of the fluorescence intensity of single exocytic events during 10 s from CgA-or CgAΔPABD-EGFP overexpressing COS-7 cells after 2 mM BaCl_2_ stimulation. The mean ± SEM of 75 exocytosis of 5 different CgA-EGFP overexpressing cells in 3 independent experiments and the mean of 65 exocytic events of 7 different CgAΔPABD-EGFP overexpressing cells in 3 independent experiments. Data have been normalized between 0 (before exocytosis) and 1 (maximum intensity). (e) Plot representing the mean ± SEM of area under curves from CgA-EGFP or CgAΔPABD-EGFP secretion kinetic. Each point represents one exocytosis event, and one color represents one analyzed cell. ****p < 0.0001, Mann-Whitney test.

Tracking EGFP-tagged CgA proteins by TIRF-M, as it has been previously performed in HEK293 and PC12 cells using CgA-EGFP^17^, revealed that PABD deletion accelerated and completed CgA release (**Figure 3c, d**) (**Videos 3** and **4**). Indeed, area under curve analysis highlighted a significant faster secretion kinetics for CgAΔPABD (2.86 ± 0.30 to 0.65 ± 0.06 AU) (**Figure 3e**), while identical Imax values indicated comparable expression levels across condition (**Supplemental Figure 3**). These results indicate that CgA’s PABD constrains secretion dynamics following stimulation.

### CgA/PA interaction regulates individual fusion events and the subsequent catecholamine release

To further dissect the role of CgA/PA interaction at the plasma membrane during neurosecretion, catecholamine release was assessed by carbon fiber amperometry, an electrochemical approach that allows to study individual fusion events in a precise and highly sensitive manner^26^. Chromaffin cells overexpressing EGFP (**Figure 4a**), CgA-EGFP (**Figure 4b**) or CgAΔPABD-EGFP (**Figure 4c**) were stimulated with a 100 mM K^+^ solution. Overexpression of CgA-EGFP did not alter the total number of exocytic events compared to control EGFP transfected cells (15.16 ± 1.08 to 12.45 ± 0.90 spikes), consistent with previous observations^17^, whereas overexpression of CgAΔPABD-EGFP significantly decreased the total number of spikes (8.51 ± 0.57 spikes) (**Figure 4d**).

**Figure 4.**
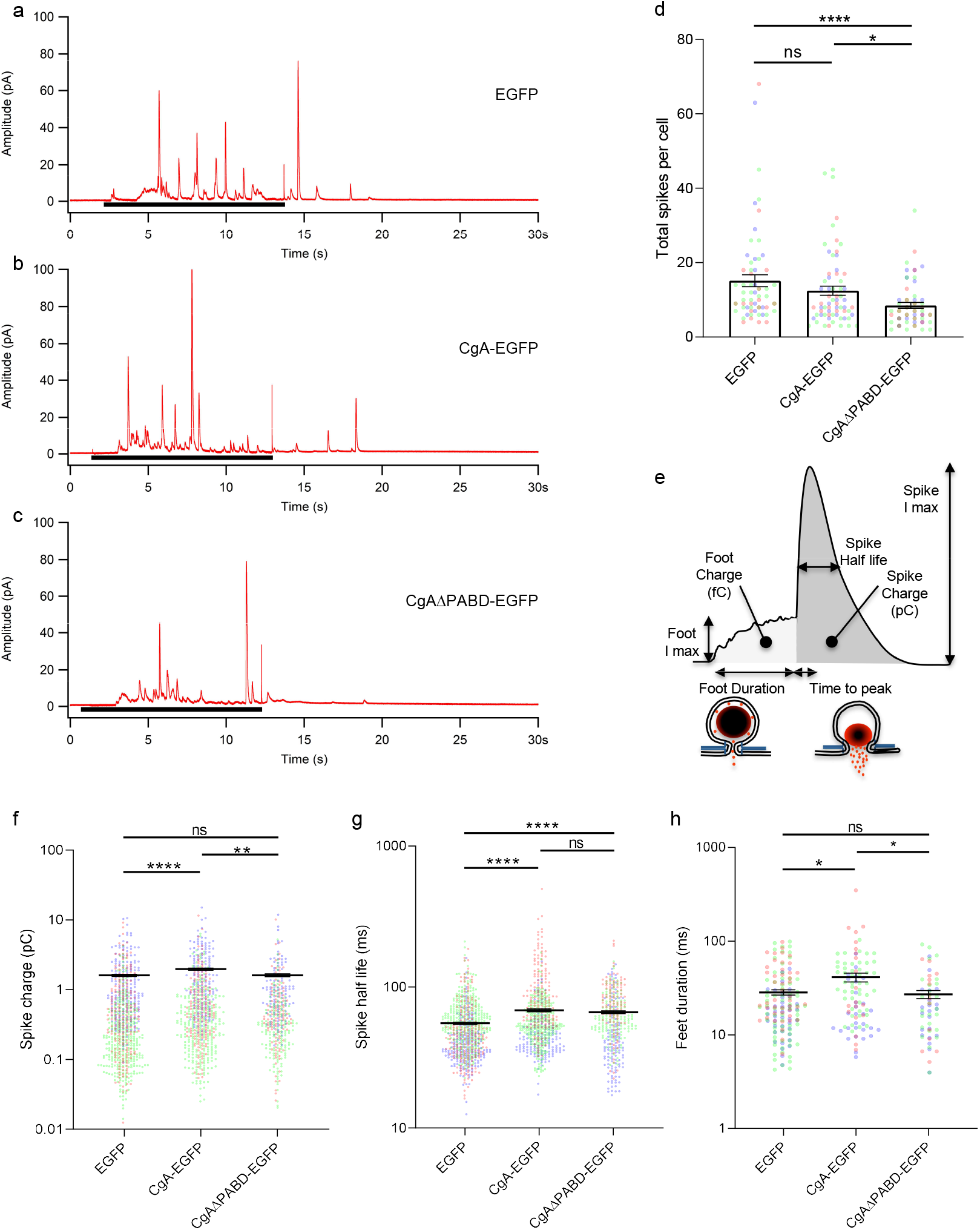
CgA/PA interaction controls catecholamine secretion. (a-c) Representation of a typical amperometrical trace obtained from chromaffin cells transfected with EGFP (a), CgA-EGFP (b) or CgAΔPABD-EGFP (c). Cells were stimulated with a puff of 100 mM depolarizing K^+^ solution indicated by the black bar under each recording. Only the first 30 s of the 60 s recordings are shown. (d) Plots representing the mean ± SEM of total spikes produced per cell (n>60 cells analyzed per condition from 3 independent cultures). (e) Description of the different individual spike parameters measured during individual spike analysis. (f-h) Plots representing the mean ± SEM of spike charge (f), spike half-life (g) (n>415 spikes analyzed per condition from 3 independent cultures) and feet duration (h) (n>60 feet analyzed per condition from 3 independent cultures) of cells overexpressing EGFP, CgA-EGFP or CgAΔPABD-EGFP. For all plots, ns: non-significant, *p<0.05, **p<0.01 ****: p<0.0001, Kruskall-Wallis followed by Dunnett’s multiple comparisons test. Each dot on the plots represents an individual measure from a cell, spike or feet. Dots are colored according to experimental repeat.

Analysis of individual fusion parameters (**Figure 4e**; **Supplemental Table 1**) revealed that CgA-EGFP increased spike charge (1.97 ± 0.05 pC) relative to both EGFP and CgAΔPABD-EGFP (1.61 ± 0.04 and 1.61 ± 0.05 pC, respectively) (**Figure 4f**), indicating enhanced granular CA loading dependent on CgA/PA interaction. CgA overexpression also slowed down pore expansion, increasing spike half-life (68.33 ± 1.02 ms) compared to control (55.31 ± 0.69 ms), with a similar effect seen for CgAΔPABD-EGFP (66.28 ± 1.21 ms) (**Figure 4g**), consistent with potential membrane independent influences on granular matrix expansion. The same effects were also observed when analyzing spike time to peak (**Supplemental Figure 4a**). Hence, the increase of spike charge and decrease of its release dynamic impaired spike maximal intensity only for CgAΔPABD-EGFP overexpression (**Supplemental Figure 4b**).

Finally, as reported previously^18^, CgA overexpression also prolonged fusion pore lifetime increasing prespike foot duration from 29.72 ± 1.96 ms (EGFP) to 41.12 ± 4.46 ms (CgA-EGFP) whereas CgAΔPABD-EGFP failed to reproduce this increase (27.04 ± 2.72 ms) (**Figure 4h**). Foot amplitude remained unchanged, leading to increased foot charge in CgA-EGFP expressing cells (**Supplemental Figure 4c, d**). Altogether these results demonstrate that CgA/PA interaction is necessary for an appropriate number of fusion events, enhances CA release, and prolongs fusion pore stability, while also revealing PA-independent CgA functions on CA release kinetics.

### CgA/PA interaction regulates exocytosis-endocytosis coupling

TEM analysis of plasma membrane sheets of stimulated chromaffin cells revealed dense vesicular structures double labelled with anti-CgA and anti-DBH antibodies (**Figure 5a**). In parallel, stimulated living chromaffin cells incubated with anti-CgA and chased for 5 min displayed increased intracellular CgA-positive vesicles (**Figure 5b, c**). Given CgA/PA interaction at the plasma membrane after exocytosis (**Figure 2d**), we tested whether these structures reflected exocytosis-endocytosis coupling.

**Figure 5.**
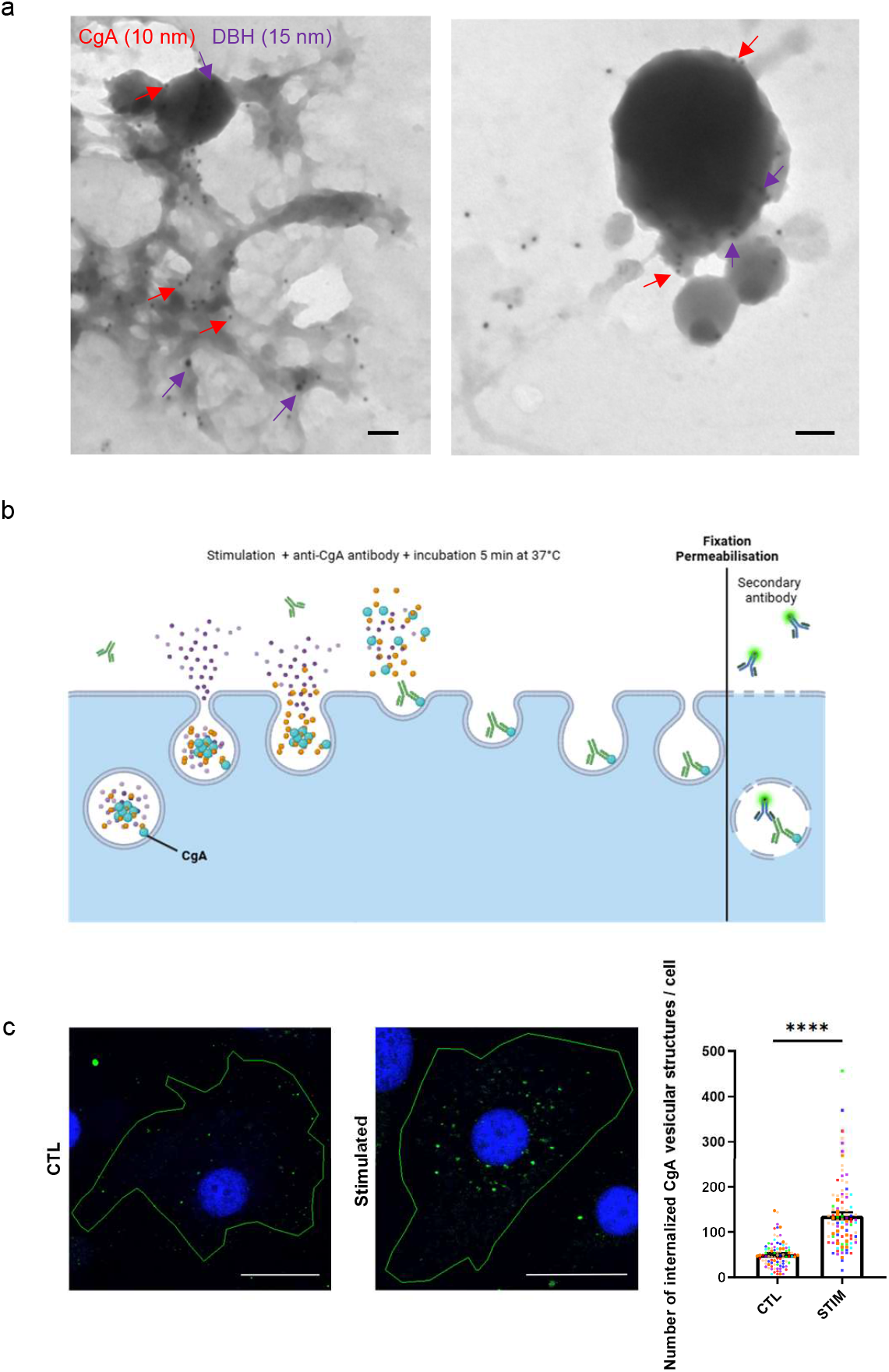
Plasma membrane-interacting CgA is internalized after regulated exocytosis. (a) Electron microscopy of membrane sheets from bovine chromaffin cells stimulated with 20 µM nicotine showing dual labelled DBH (15-nm gold beads, purple arrows) and CgA (10-nm gold beads, red arrows) vesicular structures with distinct size associated to plasma membrane. Scale bars: 100 nm. (b) Experimental protocol used for monitoring CgA internalization in living chromaffin cells. (c) Representative confocal images of chromaffin cells in resting condition (CTL) or stimulated (STIM) with 40 µM acetylcholine, incubated with anti-CgA antibody at 4°C during 45 min and then maintained at 37°C during 5 min. Cells were then fixed and processed for immunofluorescence with Alexa-488-conjugated secondary antibody to examine CgA recapture. Scale bars: 10 µm. Plot represents the quantification of endocytosed CgA-positive structures (analysis of 92 cells for the resting condition and 94 cells for the stimulated condition from 7 independent experiments). Each point is one analyzed cell and each color is an independent experiment. ****p < 0.0001, Mann-Whitney test.

Exocytic spots were labeled using anti-DBH antibody on living chromaffin cells overexpressing EGFP, CgA- or CgAΔPABD-EGFP in resting and stimulated conditions and internalization was quantified after different chase times (**Figure 6**). In resting cells, DBH signal was comparable across conditions (**Figure 6a, c**). However, stimulation with 59 mM K^+^ for 3 min markedly increased DBH signal in EGFP (19.35 ± 1.32 AU) and CgA-EGFP transfected cells (16.56 ± 0.92 AU), while the increase was significantly lower in CgAΔPABD-EGFP transfected cells (12.57 ± 1.14 AU) (**Figure 6b, c**), consistent with the reduction in fusion events observed previously (**Figure 4d**).

**Figure 6.**
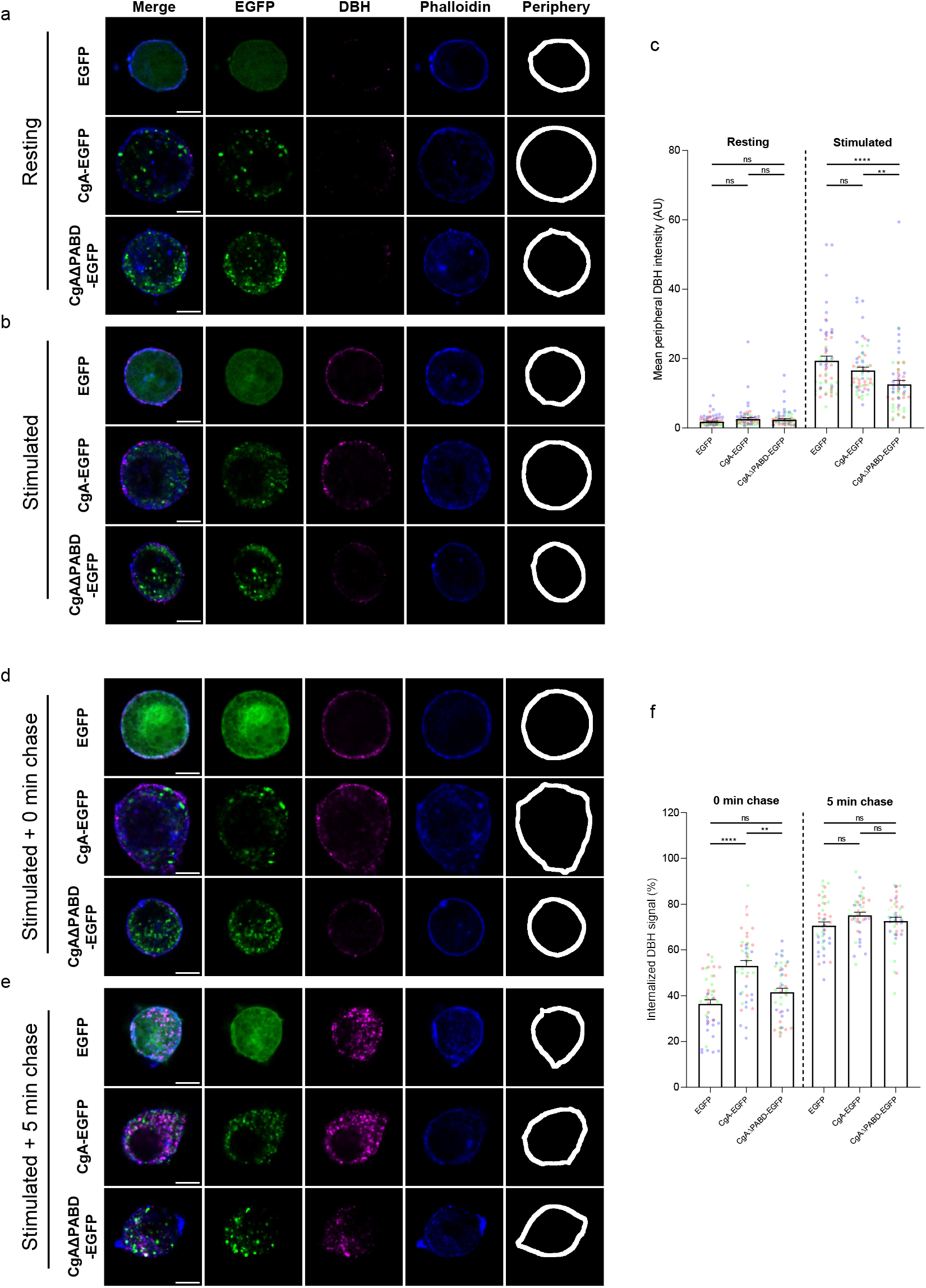
CgA/PA interaction impacts exocytic spots and their internalization. (a-b) Representative confocal images of chromaffin cells overexpressing EGFP, CgA-EGFP or CgAΔPABD-EGFP: cells were kept in resting (a) or stimulated (59 mM K^+^) (b) conditions with anti-DBH antibody for 3 min at 37°C. Cells were stained with phalloidin to segment their periphery (green: EGFP; magenta: DBH; blue: actin). (c) Plots representing the mean DBH intensity in the peripheral region (800 nm from the actin cortex) of resting and stimulated cells. Data are represented as mean ± SEM, and each dot represents a measure from a single cell, dots are also colored depending on experimental repeat (n>57 cells analyzed per condition from 3 independent cultures). ns: non-significant, **p<0.01 ****: p<0.0001, Kruskall-Wallis followed by Dunnett’s multiple comparisons test. (d-e) Representative confocal images of chromaffin cells overexpressing EGFP, CgA-EGFP or CgAΔPABD-EGFP: cells were stimulated with 59 mM K^+^ for 3 min with anti-DBH antibody and left for 0 min (d) or 5 min (e) in Locke’s solution at 37°C. (f) Quantification of the proportion of internalized DBH signal compared to the total DBH signal in cells (at least 800 nm inwards from the cortical actin cortex). Data are represented as mean ± SEM, and each dot represents a measure from a single cell, dots are colored depending on experimental repeat (n>43 cells analyzed per condition from 3 independent cultures). ns: non-significant, **p<0.01 ****: p<0.0001, Kruskall-Wallis followed by Dunnett’s multiple comparisons test. Scale bars: 10 µm.

We next assessed DBH internalization in transfected cells after 3 min stimulation followed by different chase times (**Figure 6d, e, f**). After 0 min chase, CgA-EGFP overexpression significantly enhanced DBH internalization (53.07 ± 2.29 %) relative to EGFP and CgAΔPABD-EGFP (36.42 ± 1.85 and 41.52 ± 1.79 %, respectively) (**Figure 6d, f**), whereas this difference disappeared after 5 min chase (**Figures 6e, f**). This increase in internalization indicates that CgA promotes exocytosis-endocytosis coupling through its interaction with PA via its PABD.

## Discussion

The RSP represents a fundamental cellular pathway that relies on precise membrane dynamics to sustain exocytosis in neurosecretory cells. We previously identified a direct interaction between CgA and PA using *in vitro* models and showed that this interaction contributes to SG biogenesis at the TGN, as evidenced by the reduction of SG number and the dense-core diameter decrease upon overexpression of CgA lacking its PABD^9^. Other studies further demonstrated that CgA modulates fusion pore behavior and governs CA release^17,18^.

In the present work, we extend these findings by showing that SG membrane-associated CgA interacts with PA at the plasma membrane to fine-tuning of CA storage and release, fusion pore expansion, and exocytosis-endocytosis coupling. Using complementary approaches including FLIM-FRET, TIRF-M, TEM, confocal microscopy, and carbon fiber amperometry, we analyzed CgA/PA interaction during regulated secretion and linked it to specific functional outcomes. Endogenous CgA was detected at exocytic sites after stimulation, and FLIM-FRET demonstrated that this localization reflects direct interaction with PA through the CgA PABD, marking this the first report of protein-lipid interaction monitored by FLIM-FRET in living cells using a relevant biocompatible probe^20^. Functionally, CgA-EGFP seems to be released slowly, consistent with previous observations in HEK293 and PC12 cells^17^, whereas deletion of the PABD led to markedly faster, more complete release. This difference suggests that CgA lacking PABD behaves more like a soluble neuropeptide, similar to neuropeptide Y^27^. The slower, partial evacuation of CgA-EGFP compared with the rapid, complete discharge of CgAΔPABD-EGFP demonstrates direct control of CgA exocytosis kinetics by CgA/PA interaction.

Interestingly, while PABD deletion increased the number of CgA exocytic events in COS-7 cells, it decreased the total number of CA spikes in chromaffin cells, consistent with a previously reported role for CgA composition in SG exocytosis^18^. While chromaffin cells co-expressed several granins that are known to compensate the granulogenic function of CgA^28^, we propose that, in non-endocrine COS-7 cells, CgA overexpression drives SG biogenesis, whereas CgAΔPABD expression may generate leakage through the constitutive secretory pathway, as we previously demonstrated in this cell model in the absence of the granulogenic actor myosin 1b^29^, leading to increased CgA release in basal conditions. Alternatively, CgAΔPABD expression may also generate abnormal SG. Altered CgA/PA interaction may therefore reshape granule molecular composition, as CgA enrichment could favor PA accumulation, lipid remodeling, and SG formation. Supporting this view, CgA- and PLD1-deficient mice display fewer and morphologically altered SG in chromaffin cells^30,31^. Thus, CgA/PA interaction likely regulates not only fusion dynamics but also CA release by modulating SG amount, size, and potentially the presence of transporters such as VMAT2 involved in CA loading.

Consistent with this, carbon fiber amperometry revealed increased spike charge upon CgA-EGFP overexpression, suggesting enhanced CA storage, in agreement with previous findings^17^. Importantly, CgAΔPABD overexpression failed to increase charge and reduced maximal spike amplitude. These findings suggest that CgA/PA interaction facilitates CA loading, possibly by influencing recruitment or activity of CA transporters like VMAT2. Moreover, CgA prolonged fusion pore lifetime, an effect that is absent with CgAΔPABD, showing that CgA/PA interaction also governs pore expansion. This observation aligns with a role for polyunsaturated PA species in establishing the initial fusion pore and defining release characteristics^19,32^. We propose that CgA/PA interaction stabilizes the omega-shaped intermediate during exocytosis, thereby enabling adaptive control of neuropeptide and CA release according to physiological demand^33^. In line with previous hypotheses^18^, our findings support a hypothetical model in which CgA *(i)* interacts with the highly curved fusion pore, *(ii)* engages inner leaflet lipids of the SG membrane, and *(iii)* shapes SG composition during biogenesis, mechanisms that likely coexist in neurosecretory cells expressing CgA.

Because SG membrane components remain clustered after fusion and are selectively retrieved through compensatory endocytosis^25,34^, we further investigated the role of the CgA/PA interaction in exocytosis-endocytosis coupling. In chromaffin cells, we detected membrane associated CgA with multiple makers of exocytic spots and observed stimulation dependent CgA internalization. Moreover, following CgA-EGFP overexpression, DBH internalization is increased only during the 3 min cell stimulation, but quickly caught up after 5 min chase, this effect was not observed with the PABD deleted CgA-EGFP mutant. Hence, this suggests that CgA/PA interaction promotes coordinated retrieval. This interpretation is supported by reports that clathrin and dynamin interact with mono- and polyunsaturated PA^14,20^. Thus, CgA/PA-dependent coupling may be essential to preserve plasma membrane homeostasis, regulate secretion kinetics, and recycle SG components for subsequent rounds of fusion.

Together, our findings identify CgA as a central regulator of membrane dynamics through its interaction with PA, spanning SG biogenesis at the TGN to exocytosis-endocytosis coupling at the plasma membrane. This proteo-lipid interaction orchestrates the fine control of SG content release during neurosecretion and provides mechanistic insights into pathological contexts characterized by altered CgA levels, PA synthesis, and/or CA secretion, including neurodegenerative diseases^35,36^ and tumorigenesis^37,38^. Future work should unravel how CgA, PA and associated membrane components dynamically cooperate to regulate neurosecretion, paving the way toward novel targeted therapeutic strategies.

## Methods

### Cell culture

#### Primary culture of bovine chromaffin cells

Chromaffin cells were obtained from adrenal glands of bovines aged 6 to 24 months, collected from a local slaughterhouse (according to authorization n°1069/2009 DDPP76/SPAE). After removing the surrounding fat and connective tissue from these glands, they were rinsed with 75% ethanol and then placed in cold Dulbecco’s Modified Eagle’s Medium (DMEM, Gibco, Thermo Fisher Scientific) with a mix of antibiotics and fungicids (Anti-Anti, Gibco, Thermo Fisher Scientific). The adrenal vein was first perfused with approximately 30 mL of Hank’s Balanced Salt Solution (HBSS, Gibco, Thermo Fisher Scientific) to remove blood from the vasculature. Then, each gland was digested with an enzymatic solution containing *Clostridium histolyticum* Collagenase P (Roche), DNAse (Gibco, Thermo Fisher Scientific) and trypsin (Sigma-Aldrich) (0.75 mg/0.62 mg/22.5 U per ml respectively), for 30 min at 37°C, with stirring, before being dissected in order to separate the cortical and medullary parts. The adrenal medulla was then dissociated mechanically *via* aspirations / repressions. A last incubation of the minced adrenal medulla with the enzymatic solution was carried out for 10 min at 37°C with stirring. A further dissociation was carried out by filtration through a 1 mm sieve, followed by the suppression of the enzymatic solution by centrifuging for 8 min at 500 g. The obtained cell pellet was taken up in DMEM containing 1 % Anti-Anti and filtered through a sieve of 217 μm in diameter mesh and then through a 100 μm filter (Cell Strainers, Dutscher) to remove residual cell aggregates. Deletion of remaining red blood cells was permitted by their hypotonic bursting using cold water during 10 s, followed by immediate addition of 30 mL of DMEM supplemented with 1% Anti-Anti at 37°C, before another centrifugation of the cell suspension for 5 min at 500 g. After removing the supernatant, chromaffin cells were taken up in complete DMEM medium containing 5% Fetal Bovine Serum (FBS, Sigma-Aldrich), 2.5% sterile-filtered HyClone Donor Equine serum (GE Healthcare, Life Sciences), 4 mM L-glutamine (Gibco, Thermo Fisher Scientific) and 100 μg/mL primocin (InvivoGen), distributed in T75 flasks and incubated for at least 1 h (37°C, 5% CO_2_). This seeding allows for the isolation of chromaffin cells from other cell types of the adrenal medulla such as fibroblasts, endothelial and cortical cells. Indeed, “non-chromaffin” cells adhere much more quickly than chromaffin cells to plastic substrates^39–41^. Then, the chromaffin cells-containing fraction was recovered and centrifuged, the pellet was taken up in 10 mL of complete DMEM and cells were counted using a Neubauer cell. Chromaffin cells were then distributed at a rate of 100 000 cells per well in 24-well plates containing glass coverslips previously coated with a mixture of poly-L-Lysine (20 μg/mL, 1h) and collagen I (200 μg/mL, overnight) for immunofluorescence studies. For CgA externalization and internalization studies, 10 000 000 cells were distributed in T75 flasks previously coated with poly-L-lysine (20 μg/mL, 1 h). Cultures were maintained for 24 h at 37°C, 5% CO_2_ and then the medium was renewed with complete DMEM supplemented with a mixture of antimitotics (10 μM of cytosine arabinoside and 10 μM of 5-fluoro-cytarabine, Sigma-Aldrich). The experiments were performed between 3 and 6 days after chromaffin cell seeding. For CgA overexpressing chromaffin cells (carbon fiber amperometry and exocytosis-endocytosis coupling experiments), cells were isolated and cultured as previously described prior to nucleofection^20^.

#### Culture of COS-7 cell line

African green monkey kidney fibroblast-derived COS-7 cells (American Type Culture Collection; CRL 1651) were cultured in Dulbecco’s Modified Eagle’s Medium (DMEM, Gibco, Thermo FisherScientific) supplemented with 5% fetal bovine serum (FBS, Sigma-Aldrich), 100 U/mL penicillin, 100 µg/mL streptomycin (Gibco, Thermo FisherScientific). Cells were maintained at 37°C in 5% CO_2_. For FLIM-FRET and TIRF-M experiments, 35 000 cells were plated in 35-mm glass bottom dishes (P35G-1.5-20C; MatTek Corporation) coated with 3% poly-L-lysin (Sigma-Aldrich).

### Antibodies

For immunofluorescence, primary antibodies used were rabbit monoclonal anti-CgA (MA5-14536; Thermo Fisher Scientific) (1:150 for living cells and 1:250 for fixed cells), rabbit polyclonal anti-CgA (WE-14)^42^ (1:500), mouse monoclonal anti-VMAT2 (sc-374079, Santa Cruz Biotechnology) (1:100). Secondary antibodies used were Alexa-488-conjugated donkey anti-rabbit IgG; Alexa-594-conjugated donkey anti-mouse IgG (Invitrogen) (1:500). For electron microscopy, primary antibodies used were mouse monoclonal anti-DBH (clone 4F10.2, EC.1.14.17.1: DBH, Merck Millipore) (1:50) and rabbit polyclonal anti-CgA (WE-14) (1:100). Same antibodies against DBH were also used for exocytosis-endocytosis coupling (1:200) and revealed using goat anti-mouse Alexa-555 (1:1000) conjugated secondary antibodies (Invitrogen).

### Plasmids and transfection

CgA-EGFP plasmid (pCgA-EGFP-N2) was provided by M.Courel (UPMC, Paris, France). PABD deletion (pCgAΔPABD-EGFP) was performed as described previously ^9^. Concerning the pCgA- and pCgAΔPABD-pHluorin, the original expression vectors are pCgA-EGFP and pCgAΔPABD-EGFP already mentioned where the EGFP encoding sequence has been removed and replaced by a superecliptic pHluorin sequence synthetized and cloned between SacII and NotI restriction sites by Genscript. pMAX-EGFP plasmids used as a control. For pCgA-mKate2 and pCgAΔPABD-mKate2 plasmids, EGFP was removed from pCgA-EGFP and pCgAΔPABD-EGFP with the specific enzymes (NotI and SacII). Gblocks® sequence coding for mKate2 was synthesized by IDT®. Ligation of mKate2 Gblocks® sequence with plasmids were done with a 1:3 ratio with a T4 ligase to transform Top10 bacteria. Plasmid purifications were done with the Macherey-Nagel NucleoSpin Plasmid® kit. All constructs were verified by restriction enzyme digestion and DNA sequencing.

COS-7 cells have been transfected with 0.8 µg of DNA plasmids coding for the human CgA coupled with EGFP (CgA-EGFP) or mKate2 (CgA-mKate2) or for the human CgA devoid of the PABD domain and coupled with GFP (CgAΔPABD-EGFP) or mKate2 (CgAΔPABD-mkate2), and 1.6 µL of Lipofectamine 2000 (Invitrogen) per petri dishes according to the manufacturer’s protocol. Five hours after transfection, cell medium was replaced by 37°C DMEM supplemented with 5% FBS. Cells were maintained for two days at 37°C and 5% CO_2_ before experiments.

Bovine chromaffin cells were directly transfected after their isolation. 8,000,000 cells per transfection point were resuspended in a DMEM low glucose medium without antimitotics and antibiotics supplemented with 1 mM L-glutamine and 10% FBS. Nucleofection was performed by using the primary-neuron AMAXA transfection kit (Lonza) according to the manufacturer’s protocol using program X-001. 5 µg of CgA-EGFP or CgAΔPABD-EGFP and 3 µg of GFP encoding plasmid were used. After electroporation, cells were resuspended in DMEM low glucose medium without antimitotic and antibiotic supplemented with 1 mM L-glutamine and 10% FBS and dispatched in bovine fibronectin (Sigma-Aldrich) coated 12 mm coverslips with 1/8 of the total volume per coverslip for DBH internalization experiments, or 35 mm petri dishes with 20 mm glass diameter (MatTek). Following seeding, medium was supplemented after 5 h with a twice concentrated antimitotic and antibiotic medium (1:1 v/v ratio). The following day, medium was replaced with classical chromaffin cell culture medium (low glucose DMEM supplemented with 1 mM L-glutamine, 10% FBS, 10 µM fluorodesoxyuridine, 10 µM cytosine arabinoside and 100 µg/mL primocin) to remove dead cells (around 70-80% mortality rate), then cells were maintained for 24-48 h at 37°C and 5% CO_2_ before experiments.

### Immunofluorescence studies

Chromaffin cells were rinsed twice with Locke’s solution (140 mM NaCl, 4.7 mM KCl, 2.5 mM CaCl_2_, 1.2 mM KH_2_PO_4_, 1.2 mM MgSO_4_, 11 mM glucose and 15 mM HEPES, pH 7.4) then incubated for 5 min at 37°C in Locke’s solution (control) or with Locke’s solution containing 40 μM ACh (stimulated). After rinsing with Locke’s solution, living cells were incubated for 45 min at 4°C with anti-CgA in Locke’s solution containing 0.2% fatty acid free BSA (Sigma-Aldrich). Following the incubation, cells were rinsed twice with Locke’s solution and then fixed with 4% paraformaldehyde (PFA, Sigma-Aldrich) in PBS for 15 min at 4°C before two 5 min with PBS rinses. Then, cells were permeabilized and blocked for 30 min with 0.3% Triton X-100 (Thermo Fisher Scientific), in PBS containing 5% of normal donkey serum (Sigma-Aldrich) (1:50) and 1% of BSA. Cells were then incubated for 2 h, at room temperature, with anti-VMAT2 antibodies, and for 1 h with secondary antibodies. Nuclei were stained with DAPI (4′,6-diamidino-2-phenylindole, Molecular probes) (1 µ/mL). To verify the specificity of the immunoreactions, primary or secondary antibodies were substituted with PBS. Immunocytochemistry experiments were observed with an upright confocal microscope TCS-SP8 equipped with a 63X oil immersion objective (NA = 1.4; Leica, Microsystems). Alexa-488 was excited at 488 nm and observed in a 505-540 nm window. Alexa-594 was excited at 594 nm and observed in a 600-630 nm window.

### Plasma membrane sheets

To label DBH and CgA present at the surface of cells undergoing exocytosis, cells were stimulated for 5 min with 20 µM nicotine and fixed with 2% PFA in PBS during 10 min at 4°C. After washing with PBS and blocking in PBS with 1% BSA and 1% acetylated BSA 3 h at 37°C, cells were incubated with anti-CgA and anti-DBH overnight at 4°C. Then the samples were washed 6 times with PBS and incubated 3 h with 10 nm gold particle-conjugated goat anti-rabbit IgG and 15 nm gold particle-conjugated goat anti-mouse IgG (Aurion). After 6 washes with PBS, plasma membrane sheets were prepared and processed as previously described^23^. Briefly, carbon coated Formvar films on nickel electron grids were inverted onto labelled cells, pressure was applied to the grids with a cork for 20 s, then the grids were lifted so that the fragments of the upper cell surface adhered to the grid. These membrane fragments were fixed in 2.5% glutaraldehyde in PBS, post-fixed with 0.5% OsO_4_, dehydrated in a graded ethanol series, treated with hexamethyldisilazane (Sigma-Aldrich), air-dried and observed using a Hitachi 7500 transmission electron microscope.

### FLIM-FRET

COS-7 cells were transfected with plasmids coding for CgA-mKate2 or CgAΔPABD-mKate2. The media was replaced by a Dulbecco’s Modified Eagle’s Medium without phenol red (Gibco, Thermo Fisher Scientific) supplemented with 5% FBS (Sigma-Aldrich) and HEPES (Cytiva) one hour before the beginning of the experiments. Cells were stimulated with a 2 mM BaCl_2_ solution for 5 min at 37°C. After rinsing, cells were incubated with a 3x30 s sonicated solution of 3 µM ATTO647N-coupled PA 36:1 probe^20^ during 5 min at 37°C. Then, the medium was removed to avoid the cell saturation with the PA probe, and cells were observed using a Stellaris 8 inverted confocal laser scanning microscope (STELLARIS 8 FALCON, Leica Microsystems, Nanterre, France) equipped with a white light laser (WLL) (440-790 nm), hybrid detectors HyD type X (HyDX), an 86X objective (HC PL APO 86x/1,20 W CORR CS2) for living cell acquisition, a conventional scanner (600 Hz and 512x512) and Airy 1 pinhole. A full bold line Okolab chamber (Ottaviano, Italy) installed on the inverted microscope was used to keep the temperature at 37°C and CO_2_ at 5% during image acquisition. The fast integrated FLIM module, so called FAst Lifetime CONtrast (FALCON, Leica Microsystems), was used for FRET-FLIM analysis as it allows to democratize the study of molecular interactions^43^. The donor’s lifetime was measured from Phasor plot mode, after WLL activation with 6% power at 570 nm, and HyDX detector was set in a 580 nm-641 nm window in photon counting mode. FLIM module was activated for the determination of fluorescence lifetime with 1-3 lines repetition to reach a minimum of 50 000 photons per image. The mean life-time has been measured at the whole cell, as well as at regions of interest (ROI) drawn at the level of the plasma membrane and secretory granules. For classical confocal imaging, WLL was set at 6% at 570 nm and HyDX1 detector was set in a 580 nm-641 nm window, and WLL was set at 0.5% at 647 nm and HyDX4 detector was set in a 660 nm-712 nm window. As a control, the acceptor’s (PA-ATTO647N) lifetime has also been measured to make sure that it doesn’t overlap with that of the donor (**Supplemental Figure 5**). For image acquisition, appropriate zoom factor and pixel size were set in coherence with samples.

### TIRF-M

Cell medium was replaced at least 3 h before experiments by a Dulbecco’s Modified Eagle’s Medium without phenol red (Gibco, Thermo Fisher Scientific) supplemented with 5% FBS (Sigma-Aldrich) and HEPES (Cytiva). Cells were stimulated after the beginning of recording with a 2 mM BaCl_2_ solution. Cells were observed using a Leica DMI6000B inverted microscope, in a thermostat-controlled chamber at 37°C, coupled with a 100X apochromatic oil immersion lens with a large numerical aperture (1.46) (HCX PL APO; Leica), EM-CCD (Electron Multiplying Charge-Coupled Device) C9100 with a 512x512 pixels resolution, and a 488 nm laser. The TIRF angle was set to obtain an evanescent field with a 90 nm penetration depth. The acquisition speed was set at 1 image each 250 ms (4 Hz). Photobleaching controls have been carried out (**Supplemental Figure 6**). Videos were analyzed through the FIJI software with a homemade macro and data were converted and normalized with a homemade Rstudio software macro. Areas under curves have been calculated with the GraphPad Prism 8 software.

### Carbon fiber amperometry

CA secretion was quantified 48 to 72 h post electroporation. Chromaffin cells were washed twice with ascorbate free Locke’s medium (140 mM NaCl, 4.7 mM KCl, 2.5 mM CaCl_2_, 1.2 mM KH_2_PO_4_, 1.2 mM MgSO_4_, 11 mM glucose, 10 µM EDTA and 15 mM HEPES, pH adjusted to 7.5). Transfected cells were detected using their EGFP signal with a 490 nm LED illumination (CoolLED). Upon finding a transfected cell, a 5 mm carbon fiber electrode (ALA scientific instruments) was positioned next to the individual recorded cell. Redox potential of the fiber was held at +650 mV compared to an Ag/AgCl reference electrode immersed in the medium. CA secretion was induced with a 10 s puff from a Femtotips micropipette (Eppendorf) containing a 100 mM K^+^ solution (44 mM NaCl, 100 mM KCl, 2.5 mM CaCl_2_, 1.2 mM KH_2_PO_4_, 1.2 mM MgSO_4_, 11 mM glucose, 10 µM EDTA and 15 mM HEPES, pH adjusted to 7.2). Cells were recorded for 60 s with an AMU130 amplifier (Radiometer Analytical), calibrated at 5 kHz, and digitally low pass filtered at 1 kHz. Analysis of individual exocytosis events was performed on Igor (WaveMetrics) using a macro as reported previously^32^. Only cells having produces at least 2 analyzable spikes were considered. All identified spikes with an amplitude over 5 pA were visually inspected for overlap or irregular shape. Non-analyzable spikes were not considered for individual spike parameters but counted for total spikes per cell. Feet parameters were obtained by quantifying the initial inflexion of spikes following a 50 Hz filtration, only feet with an amplitude of at least 2 pA were taken into consideration.

### Exocytosis-endocytosis coupling experiments

For the study of CgA internalization, chromaffin cells were washed twice with Locke’s solution. Cells were incubated for 5 min with 37°C, 40 µM ACh solution (stimulated) or Locke’s solution. Then, cells were incubated at 4°C for 45 min with anti-CgA antibody. Cells were then washed with Locke’s solution once and incubated for 5 min at 37°C in Locke’s solution. Coverslips were fixed with a 4% PFA in PBS during 15 min. After fixation, cells were permeabilized and blocked for 25 min at 37°C using 1% BSA in PBS containing 5% of normal donkey serum (Sigma-Aldrich) and 0.3% Triton X-100 before a 1 h incubation at 37°C with a donkey anti-rabbit-Alexa-488 in 1% BSA in PBS. Nuclei were stained with DAPI (1 µg/mL). To verify the specificity of the immunoreactions, primary or secondary antibodies were substituted with PBS. Immunocytochemistry experiments were observed with an upright confocal microscope TCS-SP8 equipped with a 63X oil immersion objective (NA = 1.4; Leica, Microsystems). Alexa-488 was excited at 488 nm and observed in a 505-540 nm window. For the study of DBH internalization, transfected chromaffin cells were washed twice with Locke’s solution. Then, cells were incubated for 3 min with 37°C, 59 mM K^+^ solution (85.7 mM NaCl, 59 mM KCl, 2.5 mM CaCl_2_, 1.2 mM KH_2_PO_4_, 1.2 mM MgSO_4_, 11 mM glucose, 560 µM ascorbic acid, 10 µM EDTA and 15 mM HEPES, pH adjusted to 7.2) (stimulated) or Locke’s solution (resting) containing anti DBH antibody. Following stimulation, cells were washed once with Locke’s solution and immediately fixed with a 4°C PBS-PFA 4% solution (Electron Microscopy Sciences) for 12 min. Then cells were permeabilized for 10 min with a 0.1% Triton X-100 PBS-PFA 4% solution at room temperature, followed by five PBS washes. All samples were blocked at 37°C for 1 h using in PBS containing 3% BSA and 10% goat serum (Sigma-Aldrich) before a 1 h incubation at 37°C with a goat anti-mouse-Alexa-555 in PBS containing 3% BSA. Cortical actin was stained with a 20 min incubation of phalloidin-ATTO647N (ThermoScientific) and coverslips were mounted on microscopy slides using a Mowiol 4-88 mounting medium (MilliporeSigma).

Cells were observed using an inverted confocal SP5II equipped with a 63X oil immersion objective (NA = 1.4; Leica, Microsystems) via the LAS AF software in sequential scanning, resolution was set at 512x512 pixels, a bit rate of 8, a sixfold zoom and a pinhole of 1 Airy Unit. EGFP (transfection control, CgA-EGFP and CgAΔPABD-EGFP) was stimulated with a 490 nm emitting laser, Alexa-555 (DBH staining) was stimulated with a 561 nm laser and ATTO647N was visualized with a 633 nm laser. Images were analyzed on the ICY software with a homemade macro to detect a peripherical zone of 800 nm width from the actin cortex^44^. For external DBH signal, the mean pixel intensity was measured in the peripheral area was measured. Finally, quantification of DBH internalization was performed by calculating the proportion of internalized DBH vs the total amount of DBH per cell.

### Statistical analysis, graphic presentation and illustrations

Statistical analysis was performed on Graphpad Prism 8 software by performing non-parametric Kruskall-Wallis test or Mann-Whitney test. Graphs were produced on the same software. All plots and described data represent mean ± SEM.

Figures 1a, 2a and c, 5b and Figure 7 were produced on Biorender.com.

## ABBREVIATIONS

ACh: acetylcholine
CA: Catecholamines
CgA: Chromogranin A
DBH: Dopamine-β-Hydroxylase
DMEM: Dulbecco’s Modified Eagle’s Medium
FLIM: Fluorescence Lifetime Imaging Microscopy
FRET: Förster’s Resonance Energy Transfer
PA: Phosphatidic Acid
PABD: PA-Binding Domain
RSP: Regulated Secretory Pathway
SG: Secretory Granule
TEM: Transmission Electron Microscopy
TGN: Trans-Golgi Network
TIRF-M: Total Internal Reflection Fluorescence Microscopy
VMAT2: Vesicular MonoAmine Transporter 2

## Code availability

All the codes used for analyzing TIRF-M data are deposited in https://github.com/berarcar/CodeMacros

## Acknowledgements

This work was supported by institutional funding from INSERM, University of Rouen-Normandie, IRIB, Région Normandie, the European Regional Development Funds (ERDF DO-IT2015 program), grants from Fondation pour la Recherche Médicale (FRM) (DEI20151234424) and from the Agence Nationale pour la Recherche (ANR-19-CE44-0019) to PYR, NV, and MM-H. TF and FL are co-supported by European Union and Région Normandie. We acknowledge the municipal slaughterhouse of Haguenau (France) and the slaughterhouse of Bigard Formerie (France) for providing the bovine adrenal glands. We acknowledge the European Regional Development Fund (ERDF «7D microscopy»), the GIS IBiSA and the In Vitro Imaging Platform-Strasbourg (CNRS UAR3156). LG, MB and DS (Normandie Node), SC-G and C. Royer (Alsace Node) are members of the national infrastructure France-BioImaging supported by the French National Research Agency (ANR-10-INBS-04).

## Author contributions

T.F. conducted FLIM-FRET, TIRF-M, and confocal imaging experiments, and detailed analysis. A.W. performed carbon fiber amperometry experiments, DBH exocytosis-endocytosis coupling experiments and associated confocal imaging. F.L. and L.J. conducted bovine chromaffin cell cultures, CgA exocytosis-endocytosis coupling experiments and associated confocal imaging. L.R., C.R. and S.C-G. performed membrane sheets and associated TEM imaging. M.B., D.S. and L.Ga. supervised FLIM-FRET and TIRF-M use. B.B. and A.S. synthesized the fluorescent PA probe PA-ATTO647N used during the study. C.B. and T.F. developed the macros used to analyze TIRF-M data. D.C., J.L. and L.Gr. assisted in plasmids construction. N.C. and Y.A. contributed to funding acquisition. P-Y.R. and S.B. developed and provided the PA probe. T.F., A.W., F.L., L.R., S.C-G., M.B., L.Ga., N.V. and M.M-H. interpreted the data and drafted the manuscript. All authors discussed the results and contributed to the manuscript revision. M.M-H. formulated the original hypothesis. N.V. and M.M-H. supervised the study.

## References

1. Bader, M. F., Holz, R. W., Kumakura, K. & Vitale, N. Exocytosis: The Chromaffin Cell As a Model System. Ann. N. Y. Acad. Sci. 971, 178–183 (2002).

2. Kidd, M., Modlin, I. M., Mane, S. M., Camp, R. L. & Shapiro, M. D. Q RT-PCR Detection of Chromogranin A. Ann. Surg. 243, 273–280 (2006).

3. Laguerre, F., Anouar, Y. & Montero-Hadjadje, M. Chromogranin A in the early steps of the neurosecretory pathway. IUBMB Life 72, 524–532 (2020).

4. Bartolomucci, A. et al. The Extended Granin Family: Structure, Function, and Biomedical Implications. Endocr. Rev. https://doi.org/10.1210/er.2010-0027 (2011) doi:10.1210/er.2010-0027.

5. Montero-Hadjadje, M. et al. Chromogranins A and B and secretogranin II: Evolutionary and functional aspects. in Acta Physiologica vol. 192 309–324 (2008).

6. Elias, S. et al. Chromogranin A Induces the Biogenesis of Granules with Calcium- and Actin-Dependent Dynamics and Exocytosis in Constitutively Secreting Cells. Endocrinology 153, 4444–4456 (2012).

7. Montero-Hadjadje, M. et al. Chromogranin A promotes peptide hormone sorting to mobile granules in constitutively and regulated secreting cells. Role of conserved N- and C-terminal peptides. Journal of Biological Chemistry 284, 12420–12431 (2009).

8. Taupenot, L. et al. Identification of a novel sorting determinant for the regulated pathway in the secretory protein chromogranin A. J. Cell Sci. 115, 4827–4841 (2002).

9. Carmon, O. et al. Chromogranin A preferential interaction with Golgi phosphatidic acid induces membrane deformation and contributes to secretory granule biogenesis. The FASEB Journal 34, 6769–6790 (2020).

10. Tanguy, E. et al. Lipids implicated in the journey of a secretory granule: from biogenesis to fusion. J. Neurochem. 137, 904–912 (2016).

11. Zeniou-Meyer, M. et al. Phospholipase D1 Production of Phosphatidic Acid at the Plasma Membrane Promotes Exocytosis of Large Dense-core Granules at a Late Stage. Journal of Biological Chemistry 282, 21746–21757 (2007).

12. Fulop, T. & Smith, C. Physiological stimulation regulates the exocytic mode through calcium activation of protein kinase C in mouse chromaffin cells. Biochemical Journal 399, 111–119 (2006).

13. Fulop, T., Radabaugh, S. & Smith, C. Activity-Dependent Differential Transmitter Release in Mouse Adrenal Chromaffin Cells. The Journal of Neuroscience 25, 7324–7332 (2005).

14. Liang, K., Wei, L. & Chen, L. Exocytosis, Endocytosis, and Their Coupling in Excitable Cells. Front. Mol. Neurosci. 10, 109 (2017).

15. Yoo, S. H., You, S. H. & Huh, Y. H. Presence of syntaxin 1A in secretory granules of chromaffin cells and interaction with chromogranins A and B. FEBS Lett. 579, 222–228 (2005).

16. Machado, J. D. et al. Chromogranins A and B as Regulators of Vesicle Cargo and Exocytosis. Cell. Mol. Neurobiol. 30, 1181–1187 (2010).

17. Dominguez, N., Estevez-Herrera, J., Borges, R. & Machado, J. D. The interaction between chromogranin A and catecholamines governs exocytosis. FASEB Journal 28, 4657–4667 (2014).

18. Abbineni, P. S., Bittner, M. A., Axelrod, D. & Holz, R. W. Chromogranin A, the major lumenal protein in chromaffin granules, controls fusion pore expansion. Journal of General Physiology 151, 118–130 (2019).

19. Tanguy, E. et al. Phosphatidic acid: Mono- and poly-unsaturated forms regulate distinct stages of neuroendocrine exocytosis. Adv. Biol. Regul. 79, 100772 (2021).

20. Schlichter, A. et al. Designing New Natural-Mimetic Phosphatidic Acid: a Versatile and Innovative Synthetic Strategy for Glycerophospholipid Research. Angewandte Chemie International Edition https://doi.org/10.1002/anie.202510412 (2025) doi:10.1002/anie.202510412.

21. Liput, D. J., Nguyen, T. A., Augustin, S. M., Lee, J. O. & Vogel, S. S. A Guide to Fluorescence Lifetime Microscopy and Förster’s Resonance Energy Transfer in Neuroscience. Curr. Protoc. Neurosci. 94, (2020).

22. Ceridono, M. et al. Selective recapture of secretory granule components after full collapse exocytosis in neuroendocrine chromaffin cells. Traffic 12, 72–88 (2011).

23. Gabel, M. et al. Annexin A2 Egress during Calcium-Regulated Exocytosis in Neuroendocrine Cells. Cells 9, (2020).

24. Umbrecht-Jenck, E. et al. S100A10-Mediated Translocation of Annexin-A2 to SNARE Proteins in Adrenergic Chromaffin Cells Undergoing Exocytosis. Traffic 11, 958–971 (2010).

25. Bittner, M. A., Aikman, R. L. & Holz, R. W. A Nibbling Mechanism for Clathrin-mediated Retrieval of Secretory Granule Membrane after Exocytosis. J. Biol. Chem. 288, 9177 (2013).

26. Poëa-Guyon, S. et al. The V-ATPase membrane domain is a sensor of granular pH that controls the exocytotic machinery. Journal of Cell Biology 203, 283–298 (2013).

27. Tsuboi, T., Ravier, M. A., Parton, L. E. & Rutter, G. A. Sustained Exposure to High Glucose Concentrations Modifies Glucose Signaling and the Mechanics of Secretory Vesicle Fusion in Primary Rat Pancreatic β-Cells. Diabetes 55, 1057–1065 (2006).

28. Montesinos, M. S. et al. The Crucial Role of Chromogranins in Storage and Exocytosis Revealed Using Chromaffin Cells from Chromogranin A Null Mouse. The Journal of Neuroscience 28, 3350–3358 (2008).

29. Delestre-Delacour, C. et al. Myosin 1b and F-actin are involved in the control of secretory granule biogenesis. Sci. Rep. 7, 5172 (2017).

30. Pasqua, T. et al. Impact of Chromogranin A deficiency on catecholamine storage, catecholamine granule morphology and chromaffin cell energy metabolism in vivo. Cell Tissue Res. 363, 693 (2016).

31. Tanguy, E. et al. Phospholipase D1-generated phosphatidic acid modulates secretory granule trafficking from biogenesis to compensatory endocytosis in neuroendocrine cells. Adv. Biol. Regul. 83, (2022).

32. Tanguy, E. et al. Mono- and Poly-unsaturated Phosphatidic Acid Regulate Distinct Steps of Regulated Exocytosis in Neuroendocrine Cells. Cell Rep. 32, (2020).

33. Wu, L. G. & Chan, C. Y. Membrane transformations of fusion and budding. Nat. Commun. 15, (2024).

34. Gasman, S. & Vitale, N. Lipid remodelling in neuroendocrine secretion. Biol. Cell 109, 381–390 (2017).

35. Jati, S. et al. Chromogranin A deficiency attenuates tauopathy by altering epinephrine– alpha-adrenergic receptor signaling in PS19 mice. Nature Communications 16, (2025).

36. Tanguy, E., Wang, Q. & Vitale, N. Role of Phospholipase D-Derived Phosphatidic Acid in Regulated Exocytosis and Neurological Disease. in Handbook of Experimental Pharmacology vol. 259 115–130 (Springer Science and Business Media Deutschland GmbH, 2018).

37. Gomez-Cambronero, J. Phospholipase D in cell signaling: From a myriad of cell functions to cancer growth and metastasis. Journal of Biological Chemistry 289, 22557–22566 (2014).

38. Houy, S. et al. Dysfunction of calcium-regulated exocytosis at a single-cell level causes catecholamine hypersecretion in patients with pheochromocytoma. Cancer Lett. 543, (2022).

39. Anouar, Y., MacArthur, L., Cohen, J., Iacangelo, A. L. & Eiden, L. E. Identification of a TPA-responsive element mediating preferential transactivation of the galanin gene promoter in chromaffin cells. Journal of Biological Chemistry 269, 6823–6831 (1994).

40. Waymire, J. C. et al. Bovine adrenal chromaffin cells: high-yield purification and viability in suspension culture. J. Neurosci. Methods 7, 329–351 (1983).

41. Ait-Ali, D. et al. Tumor necrosis factor (TNF)-α persistently activates nuclear factor-κB signaling through the type 2 TNF receptor in chromaffin cells: Implications for long-term regulation of neuropeptide gene expression in inflammation. Endocrinology 149, 2840–2852 (2008).

42. Montero-Hadjadje, M. et al. Localization and characterization of evolutionarily conserved chromogranin A-derived peptides in the rat and human pituitary and adrenal glands. Cell Tissue Res. 310, 223–236 (2002).

43. Alvarez, L. A. J. et al. SP8 FALCON: a novel concept in fluorescence lifetime imaging enabling video-rate confocal FLIM. Nat. Methods 20, 2–4 (2019).

44. Ceridono, M., Chasserot-Golaz, S., Vitale, N., Gasman, S. & Ory, S. Measurements of Compensatory Endocytosis by Antibody Internalization and Quantification of Endocytic Vesicle Distribution in Adrenal Chromaffin Cells. in 43–51 (2021). doi:10.1007/978-1-0716-1044-2_3.

