## supplemental figures table for "Chromogranin A regulates the dynamics of neurosecretion through its interaction with phosphatidic acid"

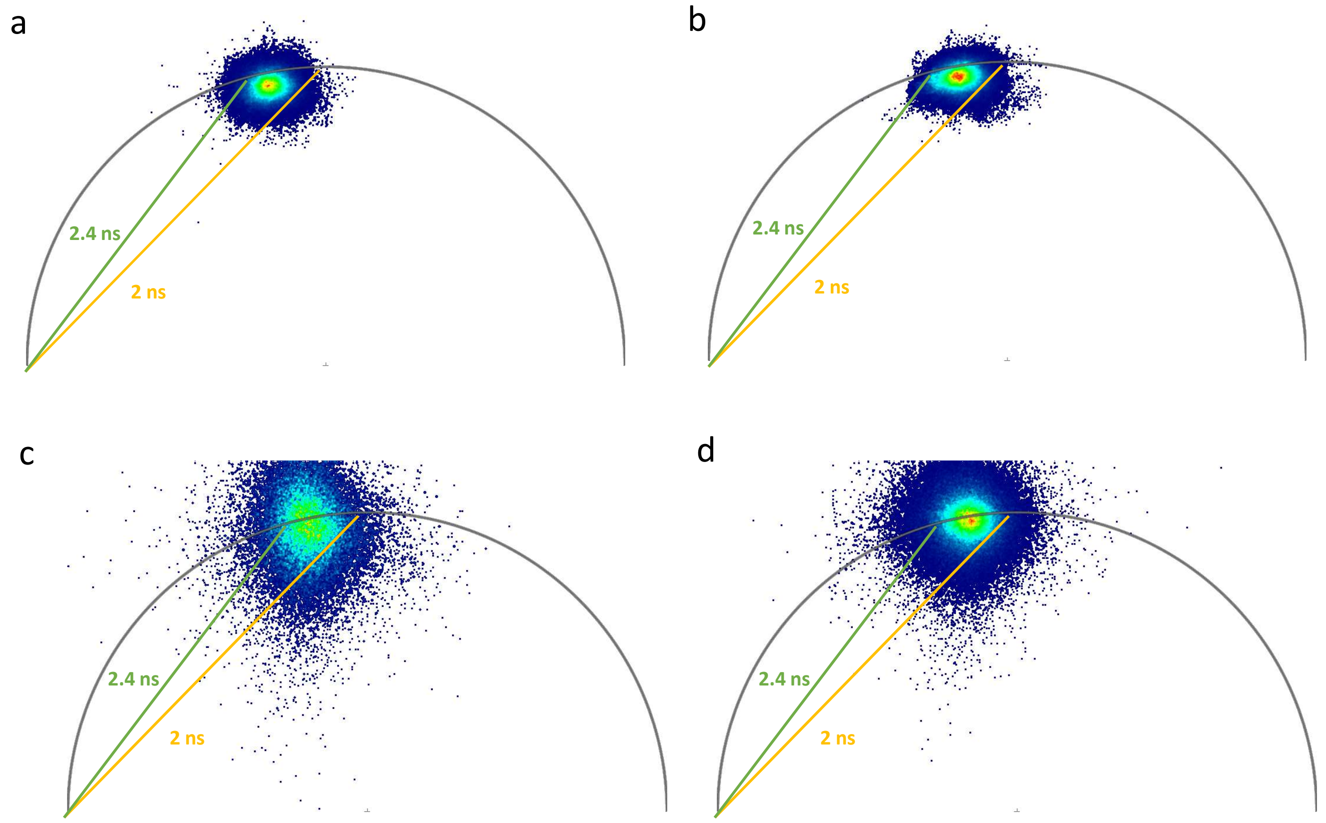


**Supplemental Figure 1: Phasor plots of CgA- and CgAΔPABD-mKate2 lifetime measured in living COS-7 cells.** (a) Phasor plot of CgA-mKate2 lifetime in stimulated living COS-7 cells overexpressing CgA-mKate2. (b) Phasor plot of CgAΔPABD-mKate2 lifetime in stimulated living COS-7 cells overexpressing CgAΔPABD-mKate2. (c) Phasor plot of CgA-mKate2 lifetime in non-stimulated living COS-7 cells overexpressing CgA-mKate2 after a 5 min incubation with 3 μM PA-ATTO647N probe. (d) Phasor plot of CgAΔPABD-mKate2 lifetime in non-stimulated living COS-7 cells overexpressing CgAΔPABD-mKate2 after a 5 min incubation with 3 μM PA-ATTO647N probe.

**Supplemental Figure 1**

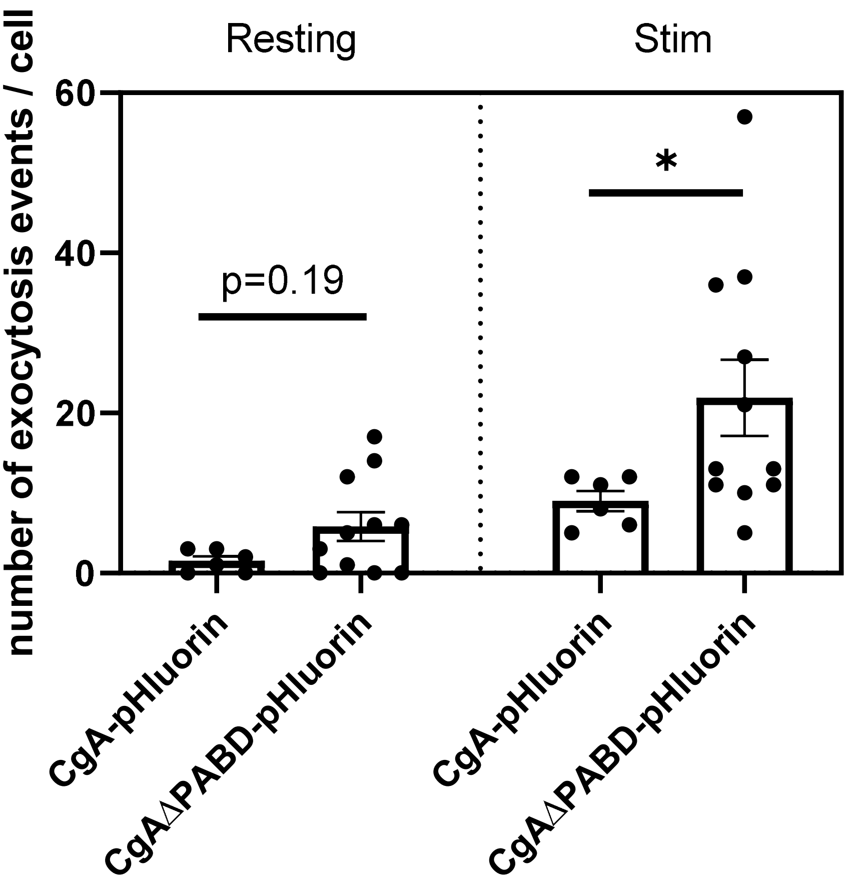


**Supplemental Figure 2: Quantification of total exocytic events before and after stimulation.** Number of exocytic events detected before (resting) and after (stim) cell stimulation with a 2 mM $BaCl_2$ solution. n = 6 and 11 cells. Mann Whitney test, * $p<0,05$.

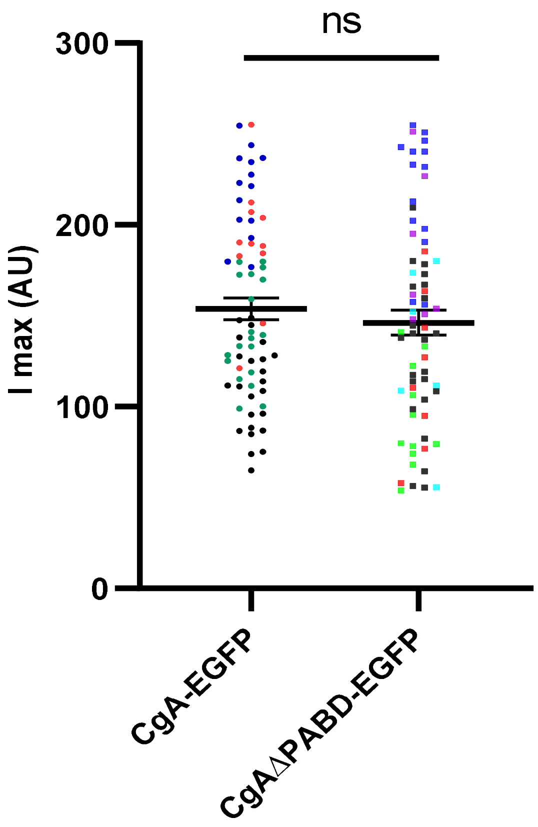


**Supplemental Figure 3: Maximal fluorescence intensities measured during exocytosis events.** Plot representing the maximum intensity of fluorescence during the exocytosis flash. 4 and 6 cells were analyzed in at least 3 independent experiments respectively for CgA and CgAΔPABD. Each point represents one exocytosis event, and one color represents one analyzed cell. AU : Arbitrary Unit. ns: non-significant, Mann-Whitney test.

**Supplemental Figure 3**

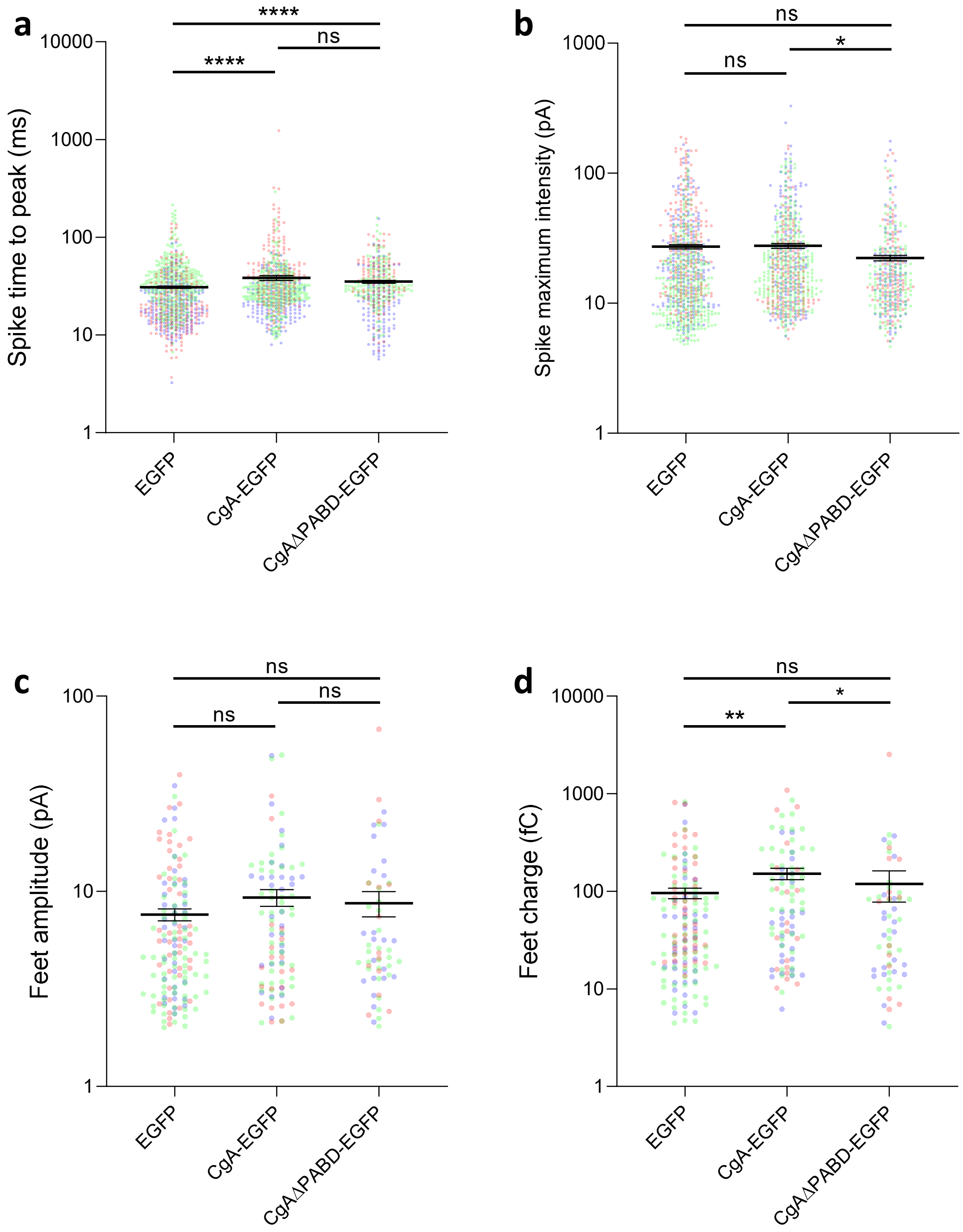


**Supplemental Figure 4: Additional quantified individual spike and feet parameters.** (a-d) Additional parameter that were quantified in the carbon fiber amperometry experiment presented on **Figure 4** of cells overexpressing EGFP, CgA-EGFP or CgAΔPABD-EGFP. Plots represent the mean ± SEM of individual spike time to peak (a), spike amplitude (b), feet amplitude (c) and feet charge (d). For all plots, ns: non-significant, *p<0.05, **p<0.01 and ****: p<0.0001, Kruskall-Wallis followed by Dunnett's multiple comparisons test. Each dot represents an individual measure from an individual spike or feet. (for a and b: n>415 spikes analyzed per condition from 3 independent cultures; for c and d: n>60 feet analyzed per condition from 3 independent cultures).

**Supplemental Figure 4**

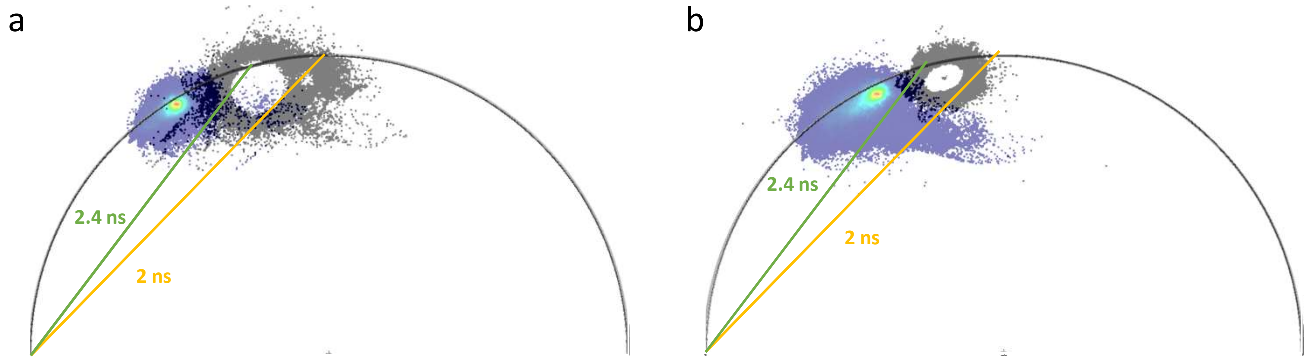


**Supplemental Figure 5: PA-ATTO647N lifetime phasor plots in living COS-7 cells.** (a) Phasor plot of PA-ATTO647N (blue) and CgA-mKate2 (grey) lifetimes in $BaCl_2$ stimulated living COS-7 cells overexpressing CgA-mKate2 after an incubation with 3 µM PA-ATTO647N probe. (b) Phasor plot of PA-ATTO647N (blue) and CgAΔPABD-mKate2 (grey) lifetimes in $BaCl_2$ stimulated living COS-7 cells overexpressing CgAΔPABD-mKate2 after an incubation with 3 µM PA-ATTO647N probe.

**Supplemental Figure 5**

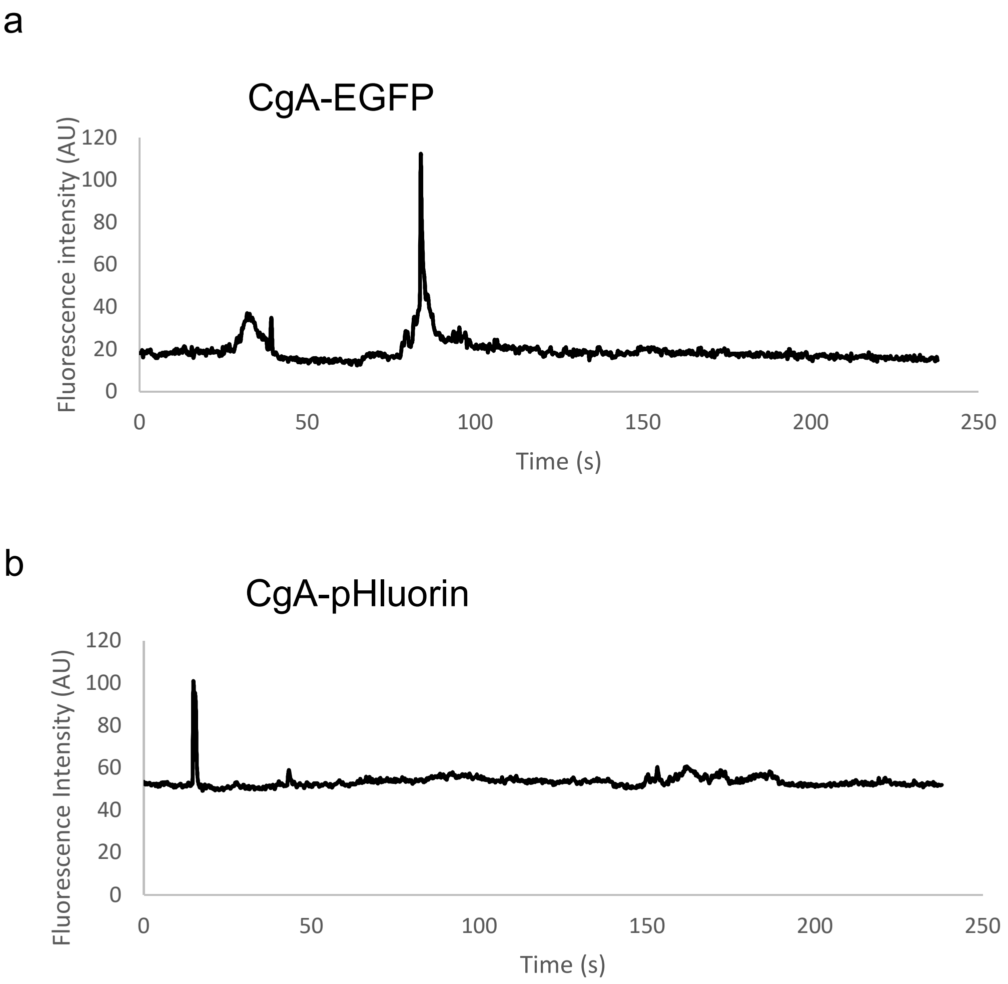


**Supplemental Figure 6: Fluorescence intensities of EGF and pHluorin during TIRF-M observations.** (a) Representative fluorescence intensity of a secretory granule containing CgA-EGFP during recording without normalization. (b) Representative fluorescence intensity of a secretory granule containing CgA-pHluorin during recording without normalization.

| | EGFP | CgA-EGFP | CgAΔPABD-EGFP |
|---|---|---|---|
| Analyzed cells | 63 | 66 | 61 |
| Spikes per cell | 15.16 ± 0.96 | 12.46 ± 0.90 | 8.51 ± 0.57 |
| Analyzed spikes per cell | 12.62 ± 0.96 | 10.17 ± 0.76 | 6.82 ± 0.47 |
| Total analyzed spikes | 795 | 671 | 416 |
| Imax (pA) | 27.21 ± 0.68 | 27.60 ± 0.70 | 22.25 ± 0.65 |
| Charge (pC) | 1.61 ± 0.04 | 1.97 ± 0.05 | 1.61 ± 0.05 |
| $T_{1/2}$ (ms) | 55.31 ± 0.69 | 68.33 ± 1.02 | 66.28 ± 1.21 |
| $T_{peak}$ (ms) | 30.83 ± 0.54 | 38.37 ± 0.77 | 35.10 ± 0.77 |
| Analyzed prespike feet | 146 | 98 | 61 |
| Feet charge (fC) | 97.35 ± 11.75 | 151.63 ± 20.04 | 119.54 ± 42.03 |
| Feet duration (ms) | 29.72 ± 1.96 | 41.12 ± 4.46 | 27.04 ± 2.72 |
| Feet amplitude (pA) | 7.96 ± 0.60 | 9.29 ± 0.93 | 8.68 ± 1.28 |

**Supplemental Table 1: Summary of all paramaters analyzed using carbon fiber amperometry on bovine chromaffin cells.** Table recapitulating all the parameters analyzed in figure 4 and supplemental figure 4, data are represented as mean ± SEM.
